# Synthetic Lethality Screening with Recursive Feature Machines

**DOI:** 10.1101/2023.12.03.569803

**Authors:** Cathy Cai, Adityanarayanan Radhakrishnan, Caroline Uhler

## Abstract

Synthetic lethality refers to a genetic interaction where the simultaneous perturbation of gene pairs leads to cell death. Synthetically lethal gene pairs (SL pairs) provide a potential avenue for selectively targeting cancer cells based on genetic vulnerabilities. The rise of large-scale gene perturbation screens such as the Cancer Dependency Map (DepMap) offers the opportunity to identify SL pairs automatically using machine learning. We build on a recently developed class of feature learning kernel machines known as Recursive Feature Machines (RFMs) to develop a pipeline for identifying SL pairs based on CRISPR viability data from DepMap. In particular, we first train RFMs to predict viability scores for a given CRISPR gene knockout from cell line embeddings consisting of gene expression and mutation features. After training, RFMs use a statistical operator known as average gradient outer product to provide weights for each feature indicating the importance of each feature in predicting cellular viability. We subsequently apply correlation-based filters to re-weight RFM feature importances and identify those features that are most indicative of low cellular viability. Our resulting pipeline is computationally efficient, taking under 3 minutes for analyzing all 17, 453 knockouts from DepMap for candidate SL pairs. We show that our pipeline more accurately recovers experimentally verified SL pairs than prior approaches. Moreover, our pipeline finds new candidate SL pairs, thereby opening novel avenues for identifying genetic vulnerabilities in cancer.

## Introduction

Synthetic lethality refers to the concept that simultaneous perturbation of gene pairs leads to cell death but individual perturbation does not [43]; see Fig. 1A for a schematic. The identification of synthetically lethal gene pairs provides a potential avenue for selective targeting of cancer cells based on genetic vulnerabilities and has already led to the development of therapies for specific patient subpopulations [39, 44]. The recent rise of large-scale perturbation screens such as the Cancer Dependency Map (DepMap) [7] offers an opportunity to identify novel SL pairs using machine learning. DepMap consists of a matrix of (real-valued) viability scores for each combination of 1078 cell lines and 17, 453 CRISPR gene knockouts; a subset of DepMap is visualized in Fig. 1B. To identify candidate SL pairs from such data, the goal is to find gene pairs (A,B), such that knockout of gene A induces a low viability score in cells that show a particular expression or mutation pattern for gene B. The difficulty in identifying SL pairs stems from the fact that the number of combinations of different expression and mutation patterns is huge.

**Figure 1:**
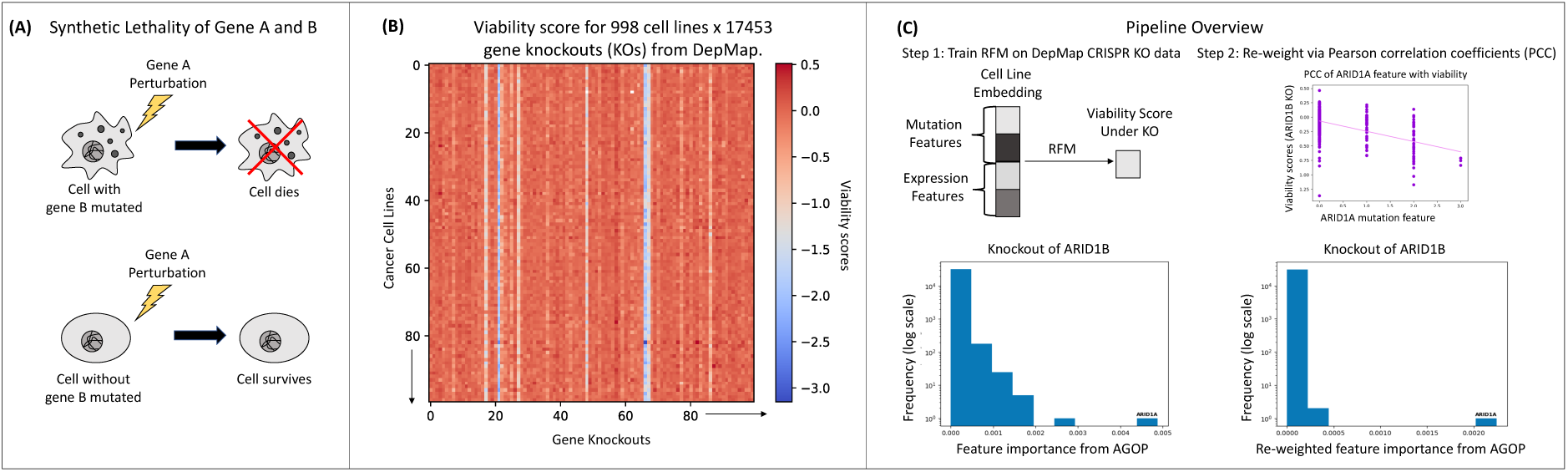
**(A)** Schematic describing the concept of synthetic lethality. Two genes (denoted A and B) form a synthetic lethality pair if the simultaneous perturbation of both genes leads to cell death but the individual perturbations do not. **(B)** A visualization of the DepMap CRISPR gene knockout data used in our pipeline. It consists of cellular viability scores for 17, 453 gene knockouts across 998 cell lines. **(C)** An overview of our pipeline. For each knockout, we train a Recursive Feature Machine (RFM) to predict the viability score from the gene expression and mutation features of a cell line. Feature importances are obtained from the trained RFM using a statistical operator known as average gradient outer product (AGOP). Given that each feature corresponds to (the expression pattern or mutation of) a gene, the genes can be ranked based on their feature importances. For example, for the knockout of ARID1B, the feature with the highest importance corresponds to the mutation of ARID1A. The second step of our pipeline re-weights the feature importances based on their Pearson correlation coefficient (PCC) with viability, thereby selecting features that are indicative of low cellular viability. The effect of such re-weighting is shown on the example of ARID1B knockout.

Various computational approaches have been developed for identifying SL pairs. Motivated by the observation that SL pairs frequently arise in paralogs (i.e., genes arising as a result of duplication) [15, 19, 20, 41], recent work trained a random forest model to predict whether a pair of paralog genes form an SL pair based on 22 hand-crafted features [18]. As such, the trained model is limited by the number of available features for prediction. Another line of work formulated SL pair identification as a link-prediction problem with links existing between SL gene pairs [4, 9, 29, 33]. This approach requires SL pairs as training data, but only few experimentally validated SL pairs are known. To overcome this limitation, these works rely on non-experimentally verified SL pairs from SynLethDB [26] or protein-protein interaction databases to augment the training set, which may lead to many false positive results. An alternative approach [55] built a predictor for the confidence of SL interactions using knockdown data from the Connectivity Map [52]; namely, it trained a neural network to map from a pair of expression vectors, corresponding to expression after knockdown of each gene in a pair, to a confidence score quantifying whether the genes form an SL pair. Labeled data for these confidence scores were obtained using GEMINI [60], a computational model for gene interactions. This again could result in many false positive results. Alternatively, statistical hypothesis testing-based pipelines were developed to characterize SL pairs, and the test results were corrected and filtered through hand-crafted criteria to balance the number of false positives and false negatives [34, 51]. Overcoming the need for any labeled SL pair data or hand-crafted statistical criteria, recent work [3] formulated the problem of SL pair screening as a feature learning task where the goal is to identify the genomic features that are most predictive of low viability under a gene knockout. This feature learning approach is the one used by the “best model” listed in the DepMap portal. Current feature learning approaches have been limited to utilizing random forests, since these simple machine learning models have been the only non-linear models that could output feature importances. If instead we could identify the features learned by state-of-the-art machine learning models, we may be better powered to find SL pairs.

In this work, we present a computationally efficient pipeline for SL pair screening by leveraging a recently developed class of feature learning methods known as Recursive Feature Machines (RFMs). RFMs were introduced in [47] and identify features learned by kernel machines, a class of machine learning algorithms that have received renewed interest in machine learning due to their connection to infinitely wide neural networks [32]. RFMs identify task-relevant features using a statistical operator, known as average gradient outer product (AGOP). AGOP has been studied in the context of task-relevant dimensionality reduction [27, 53, 59], since it identifies and amplifies the directions in data for which predictions vary the most. Given the effectiveness of kernel machines on related tasks such as virtual drug screening [48, 49] and the ability of AGOP to identify task-relevant features, we propose RFMs as a natural model for SL pair screening. Our approach is as follows: for each CRIPSR gene knockout from DepMap, we train an RFM to predict the viability score for each cell line from its mutation and gene expression features, and we use the AGOP to obtain the most predictive features. Since we are interested in pairs that reduce viability, we apply Pearson correlation-based filters to re-weight the predictive features and identify those that are indicative of low viability. Fig. 1C provides an overview of our pipeline. We will show that our pipeline recovers the experimentally verified SL pairs from [18] more accurately than previous random forest based approaches including PARIS [3] and the “best model” provided by the DepMap portal. Furthermore, by analyzing the top candidate pairs identified by our model, we obtain new candidate SL pairs that were not found using prior approaches.

## Results

### Formulating SL pair screening as a single index problem

We start by providing a novel formulation of the SL pair screening problem, which will motivate our computational approach to this problem. Recall that our goal is to find candidate SL gene pairs (A, B) such that the knockout of gene A induces a low viability score in cell lines exhibiting a particular expression or mutation pattern of gene B. We formulate this mathematically as follows. Let (*X, y_g_*) ∈ ℝ*^d×n^* × R*^n^* denote training data where *X* denotes the embedding of *n* cell lines into *d* expression and mutation features and *y_g_* denotes the viability scores when knocking out gene *g*. If *g* is part of an SL pair, then there exists a one-hot vector *u^∗^* ∈ ℝ*^d^* such that *y_g_* = *h*(*X^T^ u^∗^*) for some function *h* : R*^d^* → R, and the non-zero entry in *u^∗^* indicates the gene that forms an SL pair with *g*. This mathematical model is known as a *single-index model*, which has been extensively studied in the statistical literature [27, 28, 59] and received renewed interest in machine learning [2, 6, 17, 47]. Given that the work introducing RFMs empirically demonstrated higher sample efficiency of RFMs over other models including deep neural networks in solving such single-index problems [47], RFMs are a natural approach for SL pair screening.

### Overview of our pipeline (SL-RFM)

Our pipeline is built using the publicly available cellular viability scores for all combinations of 17, 453 CRISPR gene knockouts on 998 cell lines provided by DepMap (see Data Availability). We dropped 80 cell lines for the downstream analysis to include only the ones present in The Cancer Genome Atlas (TCGA) [57]. The first step in our pipeline is to train an RFM-based predictor to map from mutation and genomic embeddings of a cell line to the corresponding viability score under a particular knockout. To build such a predictor, we encoded each cell line as a 32, 629 dimensional vector, where the first 16, 568 real-valued features reflect gene expression and the remaining 16, 061 features represent the presence of either a damaging or hotspot mutation (see Data Availability and Methods for a detailed description of the gene expression and mutation features). We note that we used only the expression and mutation features provided by DepMap that are present in TCGA. Expression features are log transformed transcripts per million (TPM) (see Methods) and mutation features are binary (either 0 or 1) representing whether the gene has a mutation (either damaging or hotspot). After training the RFM, the model returns a list of weights for each feature denoting its importance for predicting viability for a given knockout. To ensure that the top ranked features, i.e., those with highest feature importance, are indicative of *decreased* viability, we re-weight the features by multiplying them with the Pearson correlation coefficient (PCC) between the value of the feature across all cell lines and the viability scores. Lastly, motivated by the formulation of SL pair screening as a single-index model, we identify candidate SL pairs by selecting those knockouts that contain a distribution of feature importances where the top ranked feature is well separated from the remaining features. We quantify such distributions via the difference between the maximum feature importance and the mean feature importance for a given knockout (see Methods). Training details for our pipeline are presented in Methods. In particular, the RFM framework requires the choice of a kernel. We tested the commonly used Gaussian and Laplace kernels and found that the Laplace kernel consistently outperformed the Gaussian kernel in terms of predictive performance on the DepMap data (see Methods and SI Fig. 1). Thus, all results are shown using the Laplace kernel.

### Our pipeline more accurately recovers known experimentally verified SL pairs

We first analyze the performance of our pipeline on the experimentally verified SL pairs from [18] and compare the results to the following three models: (1) a Pearson correlation coefficient (PCC) baseline, which ranks features by their correlation with viability and corresponds to the last filtration step in our pipeline; (2) a state-of-the-art random forest model for SL screening (PARIS) [3]; (3) the “best model” from the DepMap portal. The models were evaluated as follows. For each SL pair (A, B) from [18], we applied each method to predict viability under knockout of gene A and to predict viability under knockout of gene B. For each prediction task, each method outputs an importance weight for each feature, which translates into a rank based on the importance weight for the feature corresponding to gene B in the first prediction task and gene A in the second prediction task. Fig. 2A shows the minimum of these ranks for each method; SI Fig. 2 contains the ranks for each gene separately. Since the DepMap portal only provides the top 10 most important genes per knockout, for fair comparison across models the rank is denoted as *>* 10 if the pair was not identified within the top 10 most important features. We observe that SL-RFM recovers all experimentally validated SL pairs from [18] as the top ranked feature with the exception of ME2/ME3, which no model was able to identify accurately. In Fig. 2A, we also indicate whether the SL pair arises as a result of expression or mutation based features. In Fig. 2B, we present the feature importances for our method for specific knockouts from the table in Fig. 2A (the feature importance distributions for all pairs are provided in SI Fig. 3). In particular, we observe that the knockouts for which SL-RFM accurately recovered the SL pairs contain distinct feature importance patterns consistent with those expected from a single index model. Namely, there is usually one important feature corresponding to the gene that forms a synthetically lethal pair with the knockout. Note that in the one case for which SL-RFM does not identify the SL pair (ME2/ME3), there is no clear separation between the top feature and the remaining features. Lastly we emphasize the computational efficiency of our method: it involves solving a least squares problem with 32, 629 features and 998 samples for each knockout, which can be parallelized for all knockouts in a straightforward manner. As a result, the time to extract the feature importances for each knockout from SL-RFM is under 3 minutes, whereas that for PARIS is roughly 13 days.

**Figure 2:**
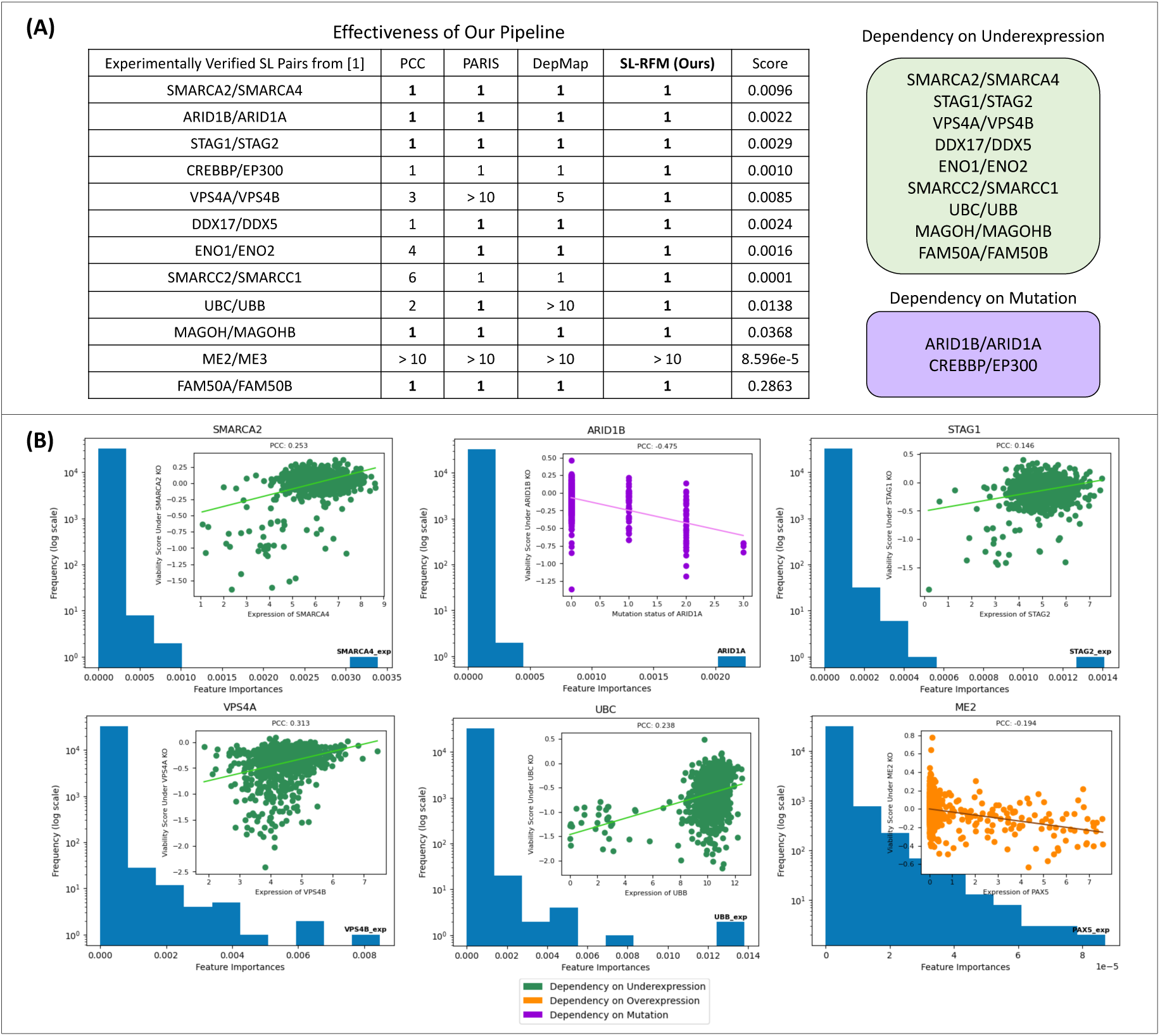
Our pipeline (denoted SL-RFM) accurately recovers experimentally verified paralog SL pairs from [18], and the feature importances for the identified SL pairs are consistent with those expected from a single-index model. **(A)** SL-RFM outperforms (1) a Pearson correlation coefficient (PCC) baseline; (2) the PARIS random forest approach from [3]; and (3) the best model from the DepMap portal described in [18]. Values in the table indicate the rank of the SL pair out of 17, 755 possible gene pairs (lower is better with a minimum value of 1). The score column quantifies the difference between maximum feature importance and mean feature importance provided by our method for the knocked out gene in the SL pair. On the right, we group SL pairs by their dependence on expression or mutation features. (**B**) Plots of the feature importance distributions for six SL pairs from [18], with the top feature labelled. Each distribution of feature importances corresponding to an SL pair identified by SL-RFM indicates a top feature that is separated from the remaining features, which is consistent with that of a single-index model. On the other hand, for the pair ME2/ME3 not identified by SL-RFM, we do not observe a clear separation between the top feature and the remaining features. The insets show how the top feature varies with viability under the given knockout.

### RFM uncovers novel candidate SL pairs

While we have thus far shown that our pipeline is able to accurately predict given experimentally verified SL pairs, the ultimate goal is to enable *de novo* identification of candidate SL pairs. Given that feature importances for known SL pairs followed our hypothesized single-index model by exhibiting a gap between the most important feature and the remaining features as shown in Fig. 2B, we used this to define a score (difference between maximum feature importance and the mean feature importance per knockout) to sort all knockouts. We selected a threshold to identify candidate SL pairs based on the elbow in the score distribution shown in SI Fig. 4. This results in 92 candidate SL pairs given by top scoring knockouts and the gene corresponding to the top feature for each knockout (see Fig. 3A as well as SI Figs. 5, 6, 7, 8 for the feature importance distributions for these SL pairs). The 4 highest scoring pairs, DDX3X/DDX3Y, EIF1AX/EIF1AY, FAM50A/FAM50B, and RPP25L/RPP25, are all experimentally verified SL pairs [18, 35, 46]. Other high scoring (and experimentally verified [46]) paralog SL pairs include CDK4/CDK6, EAF1/EAF2, and COPG1/COPG2. Notably, SL-RFM also identifies non-paralog SL pairs including MTAP/PRMT5, MTAP/WDR77, GPX4/AIFM2, and MASTL/PPP2R2A, which all have been experimentally verified [1, 5, 36]. Furthermore, we identify 327 genes that are paired with themselves, including the prominent oncogenes KRAS, BRAF, PIK3CA, which were previously identified to build SL pairs with themselves [51]. Such genes may be related to the concept of oncogene addiction [56]. In Fig. 3B, we use OncoKB [11] to validate that the majority of high scoring self-pairs are indeed oncogenes. In contrast, SI Fig. 9A shows that the majority of non-self gene pairs identified by the DepMap model that were not found by our model have not been experimentally verified. Similarly, SI Fig. 9B shows that the DepMap model proposes far more self-pairs than our model with the majority of these self-pairs not appearing in OncoKB, suggesting that the DepMap model may be producing false positives.

**Figure 3:**
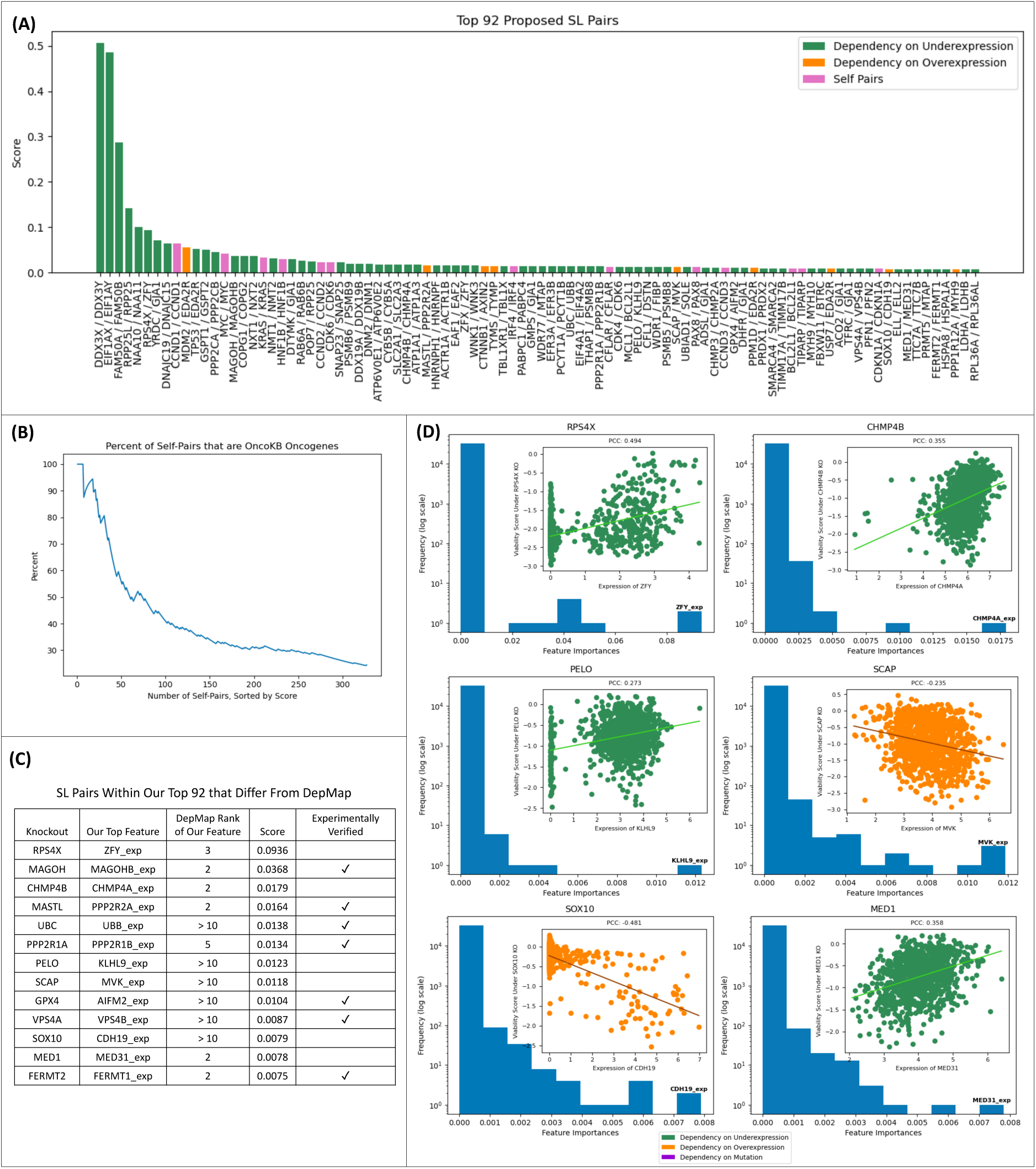
Analysis of top scoring SL pairs returned by our pipeline. **(A)** Visualization of the top 92 highest scoring SL pairs from SL-RFM and their corresponding top features, categorized by (1) if under- expression induces sensitivity to the knockout, (2) if over-expression induces sensitivity to the knockout, and (3) whether the two genes form a self pair. We note that expression features generally had more signal than mutation features, which is consistent with the findings of prior work [21]. **(B)** Validation that the majority of top scoring self-pairs identified by SL-RFM are oncogenes, based on OncoKB [11]. **(C)** A list of the SL pairs found among the top 92 highest scoring pairs from (A) for which the top feature differs from the top feature found using the best DepMap model. We observe that seven of these have been experimentally verified in prior work. (D) Distribution of feature importances for the remaining six candidate SL pairs identified from our analysis in (C) along with corresponding insets illustrating how the top feature varies with viability.

To propose new candidate SL pairs, we filtered the top 92 scoring pairs to those that did not appear using the best model from DepMap. This resulted in 13 SL pairs (see Fig. 3C). Upon further investigation, seven of these pairs identified via our pipeline have already been experimentally verified [1, 5, 18, 31, 46, 54]. The remaining six pairs identified by our pipeline, RPS4X/ZFY, CHMP4B/CHMP4A, PELO/KLHL9, SCAP/MVK, SOX10/CDH19, and MED1/MED31 all present clear characteristics of SL pairs; namely, these knockouts have a select few top features, which are associated with reduced viability (see Fig. 3D). Of these pairs, we note that only one of these knockouts (RPS4X) formed an experimentally verified SL pair when considering the top feature from the DepMap model. Indeed, DepMap identified the pair RPS4X/RPS4Y, which is a pair between an X chromosome encoded ribosomal protein gene and its Y chromosome encoded paralog [35]. SL-RFM identified RPS4Y as the second most important feature. Additionally, we note that for this knockout, nine out of the top ten ranking genes from our model are Y-linked genes, which upon their loss, are known to increase sensitivity of X-linked genes [35]. We also observe that for CHMP4B knockout and MED1 knockout, while the top features of DepMap and our model differ, the top two features of both models are CHMP4A and CHMP4C for CHMP4B knockout and MED31 and MED10 for MED1 knockout. Thus, we focus the following discussion on SCAP/MVK, PELO/KLHL9, and SOX10/CDH19 as potential novel candidate SL pairs.

MVK, or mevalonate kinase, is an essential enzyme in the mevalonate pathway, an important metabolic pathway that synthesizes isoprenoids for cellular processes [8]. Upregulation of mevalonate metabolism enhances both cancer development and the training of immunity cells [25]; thus a therapeutic challenge is to target the cancer cells without impairing the immunity cells. SCAP, or sterol regulatory element binding protein (SREBP) cleavage activating protein, regulates and chaperones genes SREBP-1 and SREBP-2, which regulate triglyceride and cholesterol levels in the body [37]. SREBP-2 appears to stimulate mevalonate metabolism [25]. *β*-catenin may bind with SREBP-2 to activate mevalonate genes and promote EMT towards an invasive cancer phenotype. On the other hand, SREBP2-dependent activation of mevalonate production seems to play a role in memory T cells, but the exact relationship is unclear [25]. Further research is needed to elucidate whether SCAP inhibition could decrease proliferation of cancer cells while enhancing immune cell response when mevalonate metabolism is upregulated.

Pelota mRNA surveillance and ribosome rescue factor (PELO) is a key gene in the cell meiotic division process. Recent work has shown that PELO and PLK1, an oncogene, bind to and degrade SMAD4, a tumor suppressor, in prostate cancer (PCa) [23]. PELO is highly expressed in PCa tissues and knockdown of PELO inhibits prostate cancer cell growth. On the other hand, kelch-like family member 9 (KLHL9) deletion is considered a driver of the mesenchymal subtype of Glioblastoma (GBM). There is a causal link between KLHL9 deletion and aberrant coactivation of transcription factors C/EBP*β*, C/EBP*α*, and STAT3, which are key regulators of this subtype of GBM [13]. While they are individually implicated in the proliferation of tumors, existing literature has not yet determined the relationship between PELO and KLHL9 in tumors. Fig. 3D shows that both the RFM features and the PCC plot indicate that knockout of PELO under low expression of KLHL9 is correlated with low cellular viability.

SOX10, or SRY-box transcription factor 10, regulates the migration of neural crest (NC)-derived cells, which are essential in embryonic development [10, 30]. Recent work has shown that CDH19, or Cadherin 19, is a direct target of SOX10, and they work together to help develop NC cells into the enteric nervous system (ENS) during embryonic development [30]. However, NC cells can also develop into melanoma. SOX10 is considered an oncogene that is heterogeneously expressed in melanoma, and its deficiency is linked to a low proliferative/high invasive phenotype [10]. While less is known about the interaction between SOX10 and CDH19 in melanoma, there is evidence that CDH19 may be in a similar oncogenic pathway as SOX10. CDH19 is widely overexpressed in melanoma, and antibodies targeting CDH19 cause tumor growth inhibition in melanoma cancer cell lines [40].

Interestingly, while the DepMap model identifies SOX10 as a self pair, SL-RFM identifies the following six genes as top features for predicting viability under SOX10 knockout: CDH19, ROPN1, EXTL1, SOX10, FGFBP2, and MIA. SI Fig. 10 shows the expression of these genes against viability under SOX10 knockout as well as the joint expression of these genes with SOX10 in TCGA. We observe a clear relationship between these genes and low viability scores for melanoma cell lines. Moreover, the oncogene SOX10 is consistently overexpressed in TCGA melanoma samples; see SI Fig. 10B. ROPN1, or Rhophilin Associated Tail Protein 1, produces a protein located in the fibrous sheath of sperm flagella [22]. Similar to CDH19, ROPN1 is an embryonic cell migration and neuronal development gene that is widely overexpressed in melanoma [16, 50]. MIA, or melanoma inhibitory activity, is used as a marker of melanoma and is a direct target gene of SOX10 [24]. Previous work has shown that SOX10 inhibition reduces MIA expression levels, which may be responsible for melanoma cell invasion [24]. There are some connections between SOX10, EXTL1, and FGFBP2 in the conjunctival epithelium. EXTL1 is a tumor suppressor that is among the top 5 most significantly upregulated factors in conjunctival melanoma [58]. FGFPB2, or Fibroblast Growth Factor Binding Protein 2, is part of the Fibroblast growth factor (FGF) system, and has been shown to be upstream of SOX9, which regulates SOX10, during the formation of ocular glands in embryonic development [14]. Reducing FGF signaling eliminated SOX10 expression, and knockout experiments in mice showed that SOX10 is essential for lacrimal gland development [14]. Another work [42] has shown that a different member of the EXT-family, EXTL2, controls FGF signaling in heparan sulfate biosynthesis. It is plausible that EXTL1 and FGFBP2 are in a pathway upstream of SOX10 that is implicated in conjunctival embryonic development and conjunctival melanoma. Overall, these experimental results suggest SOX10 as a highly attractive therapeutic target in melanoma.

### Experimental data further validates selectivity of candidate SL pairs identified by our pipeline

We first use DepMap data to demonstrate that the identified candidate SL pairs are selective across cancers. For example, if a knockout forms an SL pair based on under-expression of a gene, we expect to observe low viability in the cell lines that have low expression of the gene. On the other hand, if a knockout forms an SL pair based on over-expression of a gene, we expect to observe low viability in the cell lines that have high expression of the gene. To this end, we computed the product between the viability scores for a knockout and the expression of its suggested SL partner gene averaged across cell lines for a given cancer type. Fig. 4A shows the results for the top 92 SL pairs from our pipeline upon grouping cell lines by cancer type. This visualization highlights cancers that are predicted to be most susceptible to a given SL pair. In line with these results, somatic mutations of AXIN2, which cause over-expression of AXIN2, are associated with a higher risk of colorectal cancer and elimination of mutant CTNNB1 decreases clonogenicity of colorectal cancer cells [12, 45]. For the pair PELO/KLHL9 identified by SL- RFM in Fig. 3, which exhibited dependency on under-expression, the predicted susceptible cancers are Leukemia, Lung Cancer, and Pancreatic Cancer. For SCAP/MVK and SOX10/CDH19, which exhibited dependency on over-expression, the predicted susceptible cancers are Lung Cancer and Melanoma.

**Figure 4:**
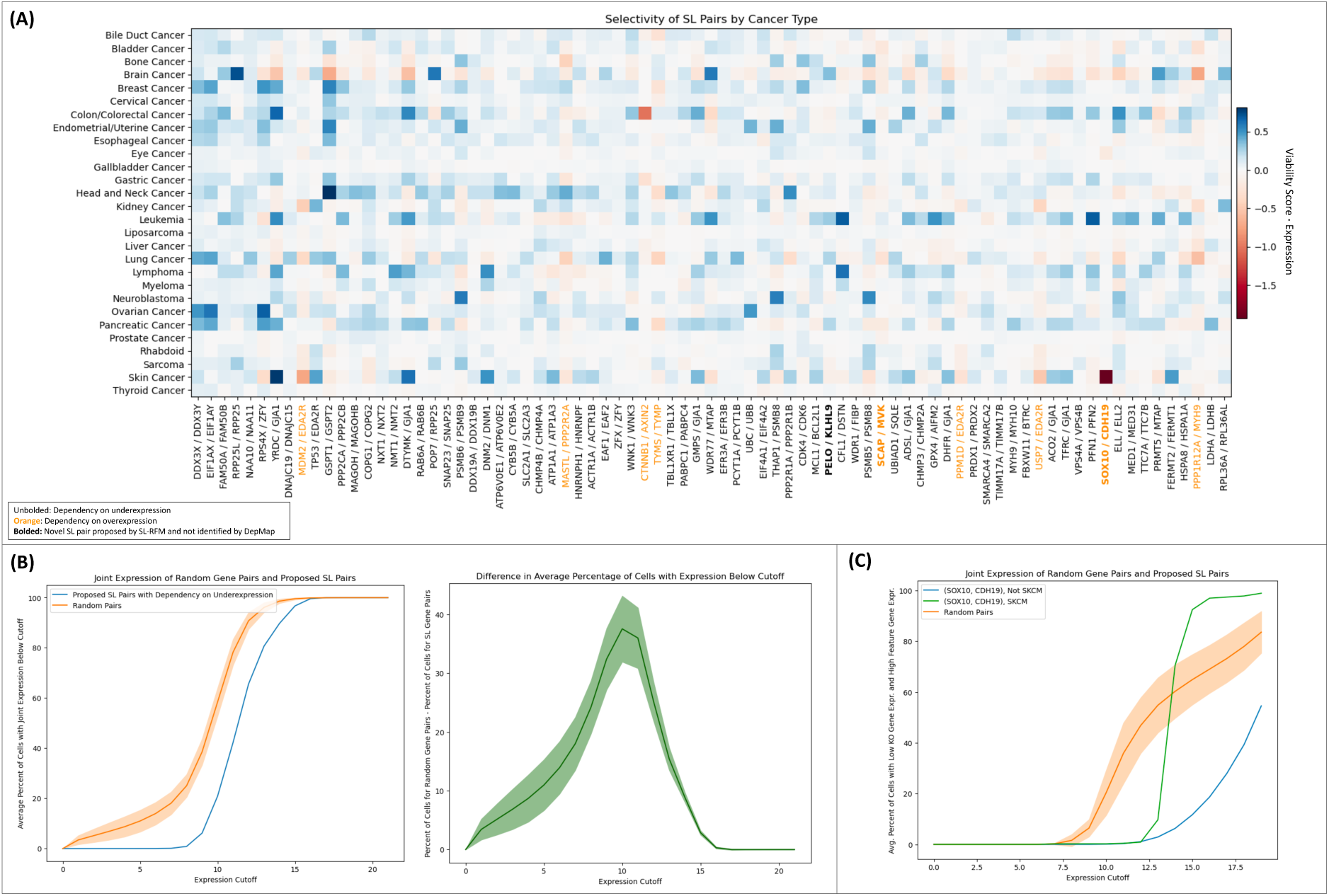
Existing experimental data further corroborates SL pairs suggested by our pipeline. **(A)** Heatmap of the predicted SL pairs and the average product between viability score of a knockout and the z-score of the expression of the top feature across DepMap cell lines aggregated by cancer type. For SL pairs with dependency on under-expression, we look for positive (blue) values. For SL pairs with dependency on over-expression, we look for negative (red) values. **(B)** Visualization validating that SL pairs with a dependency on under-expression are not simultaneously under-expressed in patient data from TCGA. Comparing the 67 out of 92 SL pairs with a dependency on under-expression that have data in TCGA (we omitted RPP25L/RPP25, COPG1/COPG2, and CHMP3/CHMP2A for this reason) to 67 randomly sampled gene pairs (sampled 10 times), we plot the average percentage of TCGA samples for which both genes have an expression below the given x-axis coordinate (error bars indicate 1 standard deviation). The curve on the right corresponds to the difference between the curves on the left. Overall, we observe that a substantial percentage of identified SL candidate pairs are never simultaneously under- expressed in patient samples. Analogous plots based on GTEx data are provided in SI Fig. 13. (C) For the proposed pair SOX10/CDH19, we plot the average percentage of TCGA samples that have expression of SOX10 below the expression cutoff, *c*, and CDH19 expression above 20 − *c* and compare with the curve for randomly sampled gene pairs (sampled 10 times) and SOX10/CDH19 in melanoma (SCKM). These curves show that there is no simultaneous under-expression of SOX10 and over-expression of CDH19 in patient samples from TCGA. Gene expression for TCGA data used in plots (B, C) was transformed via log2(normalized count + 1).

To further assess the relevance of the identified SL pairs, we used the 10, 667 samples from TCGA to confirm that the simultaneous perturbation of genes in the predicted SL pairs does not occur in patients (see Methods). Analyzing first the 67 suggested SL pairs with dependency on under-expression, Fig. 4B, compares the percentage of TCGA samples exhibiting expression below a fixed level for these SL pairs as compared to 67 randomly sampled gene pairs (averaged over 10 random draws of pairs). By definition, both curves are monotonically increasing and obtain a maximum value of 100%. Yet, we observe that early on in the graph, there is a sizable nearly 40% gap between the two curves, indicating that patient samples less often exhibit simultaneous under-expression of SL gene pairs as compared to random gene pairs. Repeating this analysis for the 9 SL pairs with dependency on over-expression (i.e., comparing the percentage of TCGA samples exhibiting low expression of gene A and high expression of gene B for these predicted SL pairs as compared to 9 randomly sampled gene pairs (averaged over 10 random draws of pairs), we observe up to 10% gap between the two curves, indicating that there are fewer patient tumor samples with the simultaneous under-expression of gene A and over-expression of gene B in the candidate SL pairs compared to random gene pairs (see SI Fig. 11). In particular, for the suggested candidate SL pairs SOX10/CDH19 and SCAP/MVK, where the dependency is on the knockout of the first gene and over-expression of the second gene, we observe that there are almost no samples with over-expression of CDH19/MVK and under-expression of SOX10/SCAP (see Fig. 4C and SI Fig. 12). Similar plots based on GTEx data [38] are provided in SI Fig. 13 and further corroborate these results. Overall, this analysis provides additional evidence that our pipeline can be used to automatically identifying potential SL pairs.

## Summary and Discussion

The identification of synthetically lethal gene pairs is a promising approach for developing targeted treatments for cancer. Large-scale perturbation screens, such as the Cancer Dependency Map (DepMap), present exciting opportunities for screening SL pairs using machine learning. Despite the availability of such data, it has been difficult to identify clinically relevant SL pairs. A promising approach has been to frame SL pair screening as a feature learning problem, where the goal is to identify, for a given knockout, the corresponding expression or mutation features most indicative of low viability. Given that random forests have been the only (nonlinear) machine learning models that provide explicit feature importances, prior work, including the “best model” from DepMap and PARIS from [3], used random forests for SL screening. In recent work, we showed that kernel machines provide a more powerful approach for related screening tasks [48, 49], suggesting that such models may provide a more powerful alternative to random forests for SL pair screening. In this work, we built a computationally efficient and effective pipeline for SL pair screening by leveraging a recently developed class of feature learning kernel machines known as Recursive Feature Machines (RFMs). The construction of our pipeline is motivated by formulating the problem of SL pair screening as the prominent single-index model studied in the statistical literature, which can be solved using an RFM. We demonstrated that our pipeline more accurately recovers experimentally verified SL pairs than the “best model” from DepMap and previous state-of-the-art approaches based on random forests. Moreover, we applied our pipeline to identify novel candidate SL pairs. We identified as top candidates PELO/KLHL9, MVK/SCAP, and SOX10/CDH19, which were not found using prior approaches but have strong supporting evidence based on experimental data from DepMap and The Cancer Genome Atlas (TCGA). In the following, we discuss implications of our results and future extensions.

### Identifying synthetically lethal gene groups

In this work, we focused on the problem of identifying gene pairs, which when simultaneously perturbed lead to cell death, by formulating it as a single-index problem. An interesting next direction is to investigate whether our pipeline can also be used to identify patterns of groups of genes that are associated with low cellular viability under a given knockout; this represents a natural extension of single-index to multi-index modeling. Another interesting extension is to consider combinatorial screens in which more than one gene is perturbed. While such settings are challenging since there is limited data on the effects of combinatorial perturbations on cells, we envision that our pipeline could serve as a useful step in determining candidate experiments for identifying novel SL gene interactions.

### Flexible and scalable approach for screening

The flexibility and computational efficiency of our pipeline, which takes less than 3 minutes to run on the full DepMap dataset on a single GPU, opens novel avenues for rapidly obtaining novel biological hypotheses of key features. In particular, while our current pipeline explores expression and mutation features, there are a number of other features including copy number, ploidy, and features from multimodal data that we could integrate to identify novel candidates of cancer vulnerabilities. We envision that integrating such features could provide novel insights into biological mechanisms associated with cancer and support the development of novel targeted treatments.

## Methods

### Overview of datasets and pre-processing

Below, we provide an outline of all datasets and data pre-processing utilized in this work.

- DepMap

**–** The **viability** dataset is a 1078 cell lines × 17,453 knockout matrix where every entry is a number denoting the viability of the cell after the gene is knocked out.
**–** The **gene expression** dataset contains a 19,194-dimensional gene expression (log2(TPM + 1)) vector for 1,408 cancer cell lines.
**–** The **damaging mutations** dataset contains a 17,257-dimensional vector for 1,702 cancer cell lines with values in [0, 1, 2], where 0 indicates no mutation, 1 indicates at least one heterozygous damaging mutation, and 2 indicates at least one homozygous damaging mutation.
**–** The **hotspot mutations** dataset contains a 450-dimensional vector for 1,702 cancer cell lines with values in [0, 1, 2], where 0 indicates no mutation, 1 indicates at least one heterozygous hotspot mutation, and 2 indicates at least one homozygous hotspot mutation.
- TCGA

**–** The TCGA dataset contains 10667 samples from patients with 36 different cancers: Acute Myeloid Leukemia (LAML), Adrenocortical Cancer (ACC), Bile Duct Cancer (CHOL), Bladder Cancer (BLCA), Breast Cancer (BRCA), Cervical Cancer (CESC), Colon Cancer (COAD), Colon and Rectal Cancer (COADREAD), TCGA Endometrioid Cancer (UCEC), Esophageal Cancer (ESCA), Glioblastoma (GBM), Head and Neck Cancer (HNSC), Kidney Chromophobe (KICH), Kidney Clear Cell Carcinoma (KIRC), Kidney Papillary Cell Carcinoma (KIRP), Large B-cell Lymphoma (DLBC), Liver Cancer (LIHC), Lower Grade Glioma (LGG), Lower Grade Glioma and Glioblastoma (GBMLGG), Lung Adenocarcinoma (LUAD), Lung Cancer (LUNG), Lung Squamous Cell Carcinoma (LUSC), Melanoma (SKCM), Mesothelioma (MESO), Ocular Melanomas (UVM), Ovarian Cancer (OV), Pancreatic Cancer (PAAD), Pheochromocytoma and Paraganglioma (PCPG), Prostate Cancer (PRAD), Rectal Cancer (READ), Sarcoma (SARC), Stomach Cancer (STAD), Testicular Cancer (TGCT), TCGA Thymoma (THYM), Thyroid Cancer (THCA), or Uterine Carcinosarcoma (UCS). These are all the cancers available on the Xena portal except for Pan-Cancer (PANCAN) and Formalin Fixed Paraffin-Embedded Pilot Phase II (FPPP). The gene expression dataset contains a 40,543-dimensional gene expression (log2(TPM + 1)) vector and mutation dataset contains a 40,543 -dimensional binary vector indicating the presence of non-silent somatic mutation.

Each cell embedding is a concatenation of the DepMap gene expression vector and DepMap mutation vector, where the mutation vector contains a 1 if the cell has a damaging or hotspot mutation and 0 otherwise. The gene expression vector is z-scored across cell lines, so the mean gene expression for each gene is 0. The cell embedding is normalized to norm 1 under the *ℓ*_2_ norm for each cell. For ease of downstream analysis, we only kept cell features that are present in both the DepMap and TCGA datasets. The final dimensions of the gene expression features and mutation features are 16,568 and 16,061, respectively. We also only kept cell lines that are present in both the viability and gene expression/mutation datasets. This leaves 998 cells. The final cell embedding is a 998 × 32, 629-dimensional matrix.

### Training details

We trained a RFM for each knockout that is trained to map cell line embeddings to viability scores for each knockout. Since the cell embeddings are the same per knockout, we trained RFMs efficiently by modeling this problem as a multi-output regression problem where the number of outputs are equal to the number of knockouts. When trained RFMs using a Laplace kernel as the base predictor, i.e., the kernel function is *K*(*x, x̃*) = exp(−*L*||*x* − *x̃*||_2_). We directly solved the kernel regression problem with *L* = 1 and performed one iteration of feature-learning through the average gradient outer product. We used ridge regularization with a coefficient of 10*^−^*^6^ to avoid numerical issues with solving exactly. If *Y* ∈ R^998^*^×^*^17453^ is the viability matrix and *X* ∈ R^998^*^×^*^32629^ is the cell embedding, then the solution to our kernel regression problem is

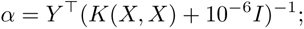

where *K*(*X, X*)*_ij_* = *K*(*x_i_, x_j_*) with *x_i_, x_j_* denoting embeddings of cell lines *i, j*. We then utilize the AGOP of the trained kernel machine to extract relevant features. In particular, for a cell line *x* and KO *k*, let the trained kernel machine be given by

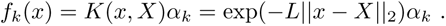

Since we are primarily interested in feature importances, we compute the average magnitude of gradient entries of *f_k_*. These are given by the diagonal of the AGOP as follows:

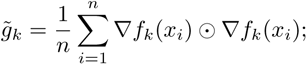

where ⊙ denotes elementwise multiplication (Hadamard product). Note that feature importance alone is insufficient for identifying SL pairs. In particular, we must ensure that the top feature induces low cellular viability. To this end, we use Pearson correlation coefficient (PCC) to additionally re-weight features based on their association with low viability. Namely, for mutation features, the presence of a mutation should be negatively correlated with the viability score. For expression features, a strong positive correlation with viability indicates that the knockout is synthetically lethal with under-expression of the feature and a strong negative correlation indicates that the knockout is synthetically lethal with over-expression of the feature. To incorporate these relationships into our pipeline, we re-weight the feature importances according to

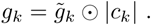

where *c_k_* is a 32, 629-dimensional vector where each entry is the PCC of the feature gene with viability under the knockout and coordinate *i* of *c_k_* is 0 if coordinate *i* represents a mutation feature and the PCC of the feature is positive. Moreover, for entries of *c_k_* corresponding to expression features, we only utilize features within 3 standard deviations of the mean to omit outliers.

### Metrics for evaluating performance

To find the best kernel to use, we benchmarked the performance of Laplacian and Gaussian kernels to predict the viability scores for each knockout. The Laplacian kernel is defined as *K*(*x, x̃*) = exp(−*L*||*x* − *x̃*||_2_) and Gaussian kernel is defined as 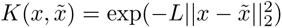 for *L >* 0. For our experiments, we used *L* = 1 for both kernels. We also benchmark predicting the mean for each knockout over cell lines. We used 5-fold cross validation for the target task and reported the metrics computed across all folds in SI. Fig. 1.

Let *C* = 998 denote the number of cell lines and *K* = 17453 denote the number of knockouts. Let *Ŷ* ∈ ℝ*^C×K^* denote the predicted viabilities for cells under each knockout generated through 5-fold cross validation. Let *Y* ∈ ℝ*^C×K^* denote the ground truth. Let 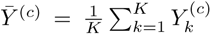 denote the average viability for cell line *i*. Let **ŷ**, **y** ∈ ℝ*^CK^* denote the vectorized versions of *Ŷ* and *Y* respectively, and 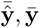 their respective means. We use the same three metrics as those considered in [48, 49]. All evaluation metrics have a maximum value of 1 and are defined below.

1. Pearson R:

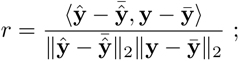

2. Mean *R*^2^:

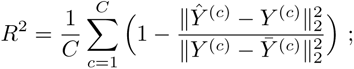

3. Mean Cosine Similarity:

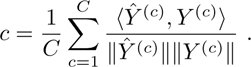

In SI Fig. 1B, we evaluate the prediction performance for computational simplicity in two specific cell lines, A549 and HUH7, using 5-fold cross validation. For each cell line, we obtain a list of knockouts sorted from most lethal to least lethal for the cell. We generate the same sorted list using predicted viability scores from the Laplacian kernel, mean over cell lines, and a random shuffle. We then plot the percentage of knockouts that overlap between these lists, indexed by the number of the most lethal knockouts. Laplacian kernel out-performs the other two benchmarks, demonstrating that SL-RFM effectively identifies the top most lethal knockouts for held out cell lines.

### Score computation from feature importances

To propose candidate SL pairs, we look for knockouts for which feature importance distributions exhibit a gap between the most important feature and other features consistent with a single-index model. We thus define the following score to automatically identify such SL pairs. Let *v_k_*be a vector of feature importances for the predictor *f_k_* trained to predict viability scores of knockout *k*. We define

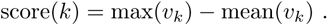

We then sort the knockouts by their scores. The knockouts with the highest scores and the genes associated with the top feature for these knockouts are proposed as candidate SL pairs.

## Data Availability

All datasets considered in this work are publicly available. The CRISPR-Cas9 viability screens (CRISPRGeneEffects.csv), cell line expression (OmicsExpressionProteinCodingGenesTPMLogp1.csv), cell line mutation data (OmicsSomaticMutations.csv), cell line (Model.csv), and DepMap feature importances (Chronos_Combined_predictability_results.csv) were downloaded from DepMap version 22Q4. The TCGA data was downloaded from the UCSC Xena datahub (tcga.xenahubs.net), and the GTEx data is Gene TPM data from GTEx Analysis V8 (GTEx_Analysis_2017-06-05_v8_RNASeQCv1. 1.9_gene_tpm.gct.gz). A list of oncogenes was obtained from OncoKB at https://www.oncokb.org/ cancer-genes.

## Code Availability

The code is available at https://github.com/uhlerlab/synthetic_lethality.

## Acknowledgements

We thank Christopher Moy and Barbara Weir for helpful discussions as well as Janssen Pharmaceuticals for their support. We also thank Todd Golub and William Sellers for their insights. A.R. was funded by an Eric and Wendy Schmidt Center fellowship as well as the George F. Carrier Postdoctoral fellowship in the School of Engineering and Applied Sciences at Harvard University. This research was supported by NCCIH/NIH (1DP2AT012345), ONR (N00014-22-1-2116), the MIT-IBM Watson AI Lab, and a Simons Investigator Award (to C.U.).

## Supporting Information

**SI Figure 1:**
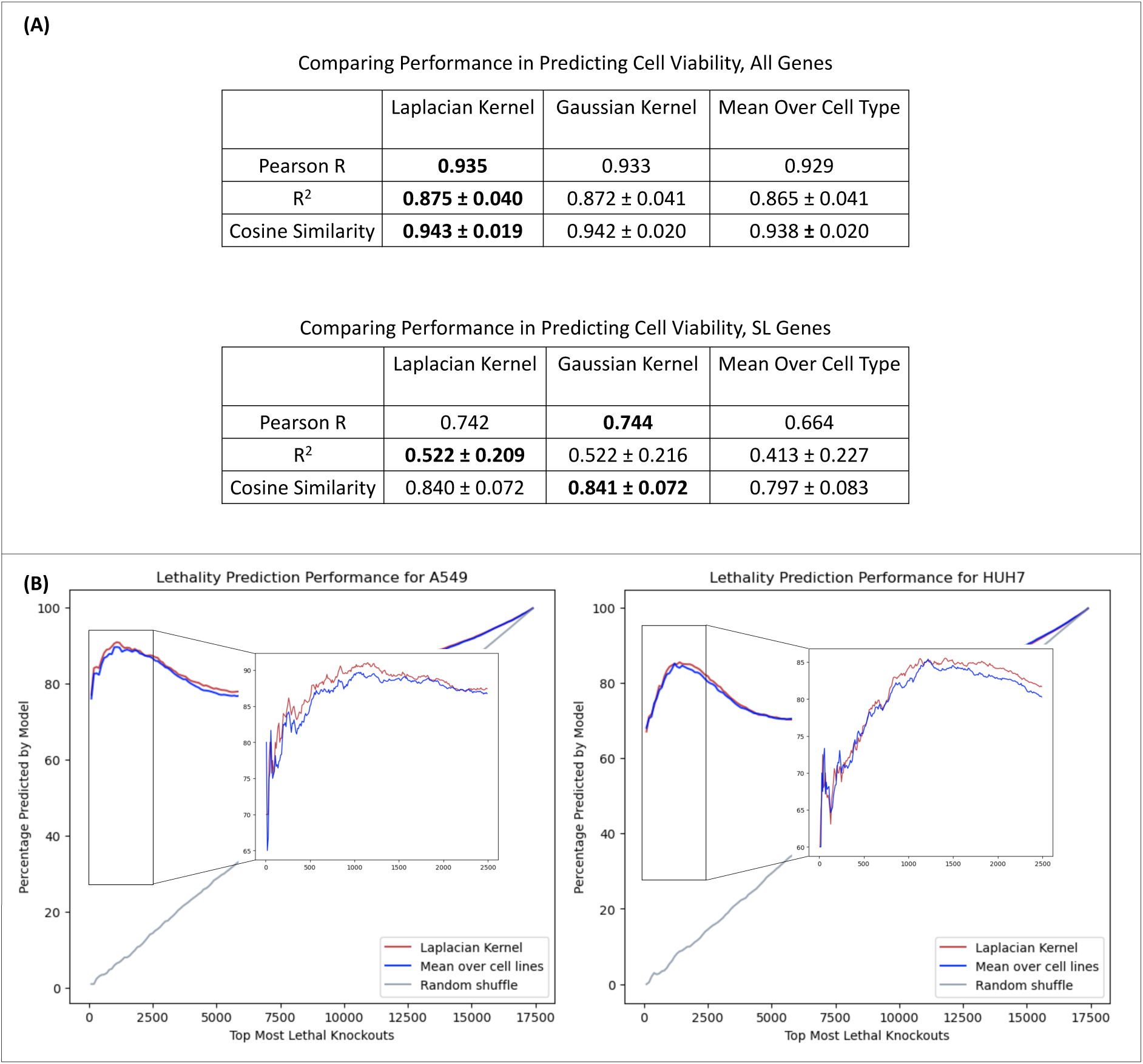
Comparison of model performance in predicting viability data. All metrics are described in Methods. **(A)** Comparison of RFM with varying base kernels (Laplace and Gaussian kernel) on predicting cell viability, where test cell lines are held out using 5-fold cross validation. **(B)** Analyzing predictions for individual cell lines from five-fold cross-validation. We compare the list of predicted knockouts and the list of ground truth knockouts sorted by viability score for sample cell lines (A549 and HUH7). We observe that RFM with the Laplace kernel base predictor outperforms both a mean over cell line benchmark and a random baseline, illustrating the effectiveness of our selected model on held-out test data.

**SI Figure 2:**
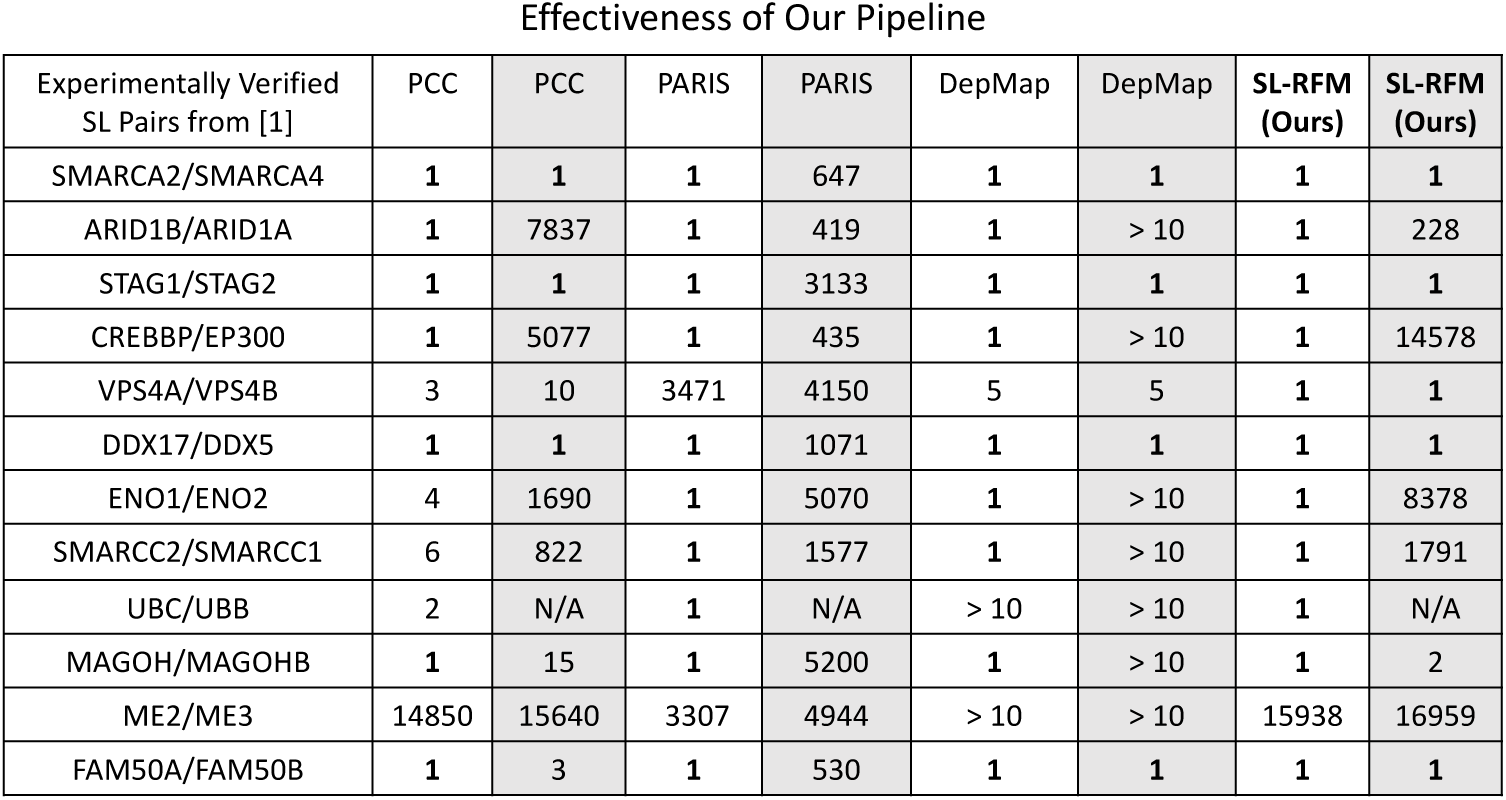
Ranks of all models from Fig. 2 when considering the first gene in the pair as the knockout (white) or the second gene in the pair as the knockout (gray). The minimum rank between white and gray columns is reported in Fig. 2. We note that DepMap only provides the top 10 most important features for prediction. If the SL pair gene does not appear among the top 10, we denote the rank as *>* 10. All other models return the full set of feature importances.

**SI Figure 3:**
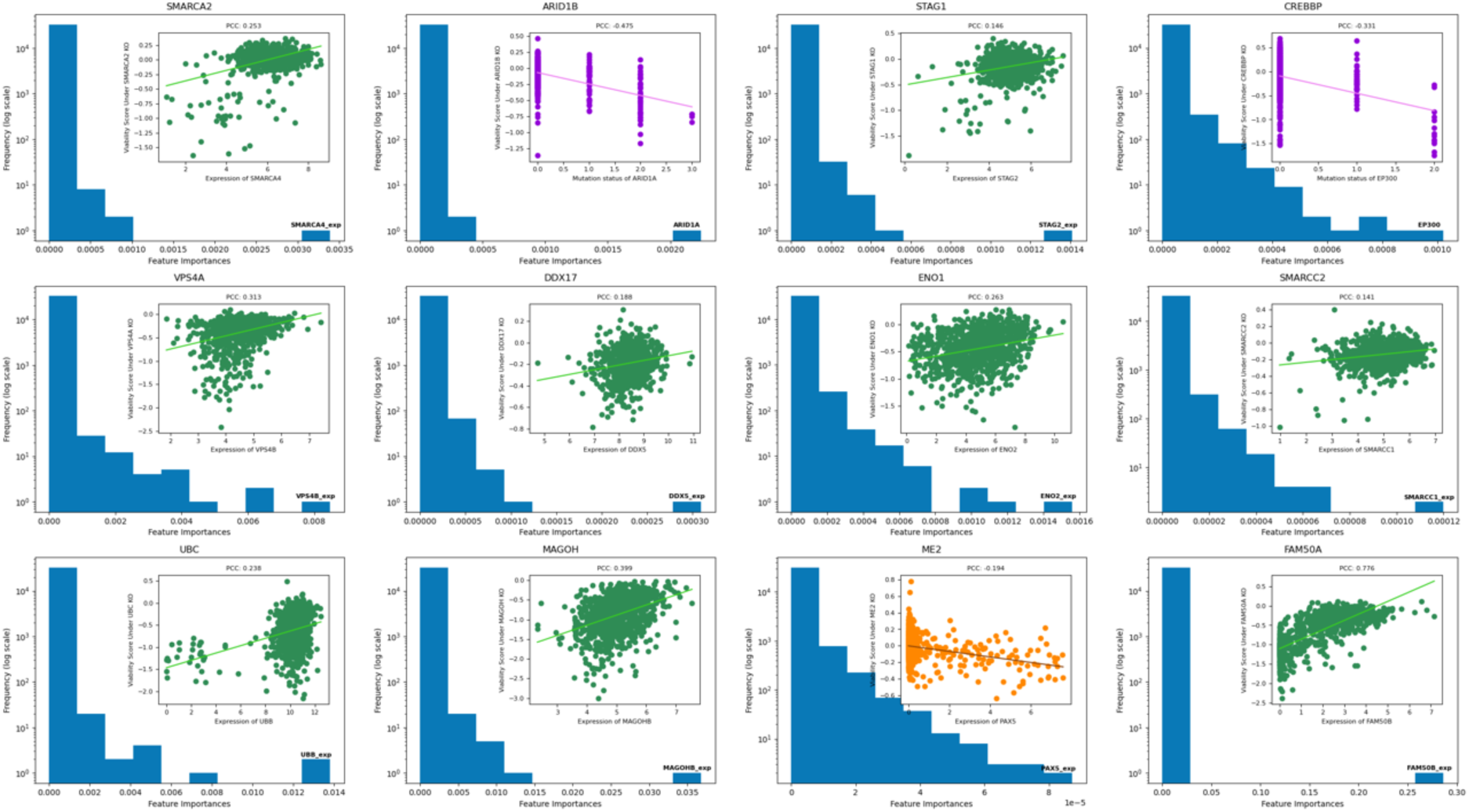
SL-RFM feature importance plots for knockouts part of SL pairs from [18].

**SI Figure 4:**
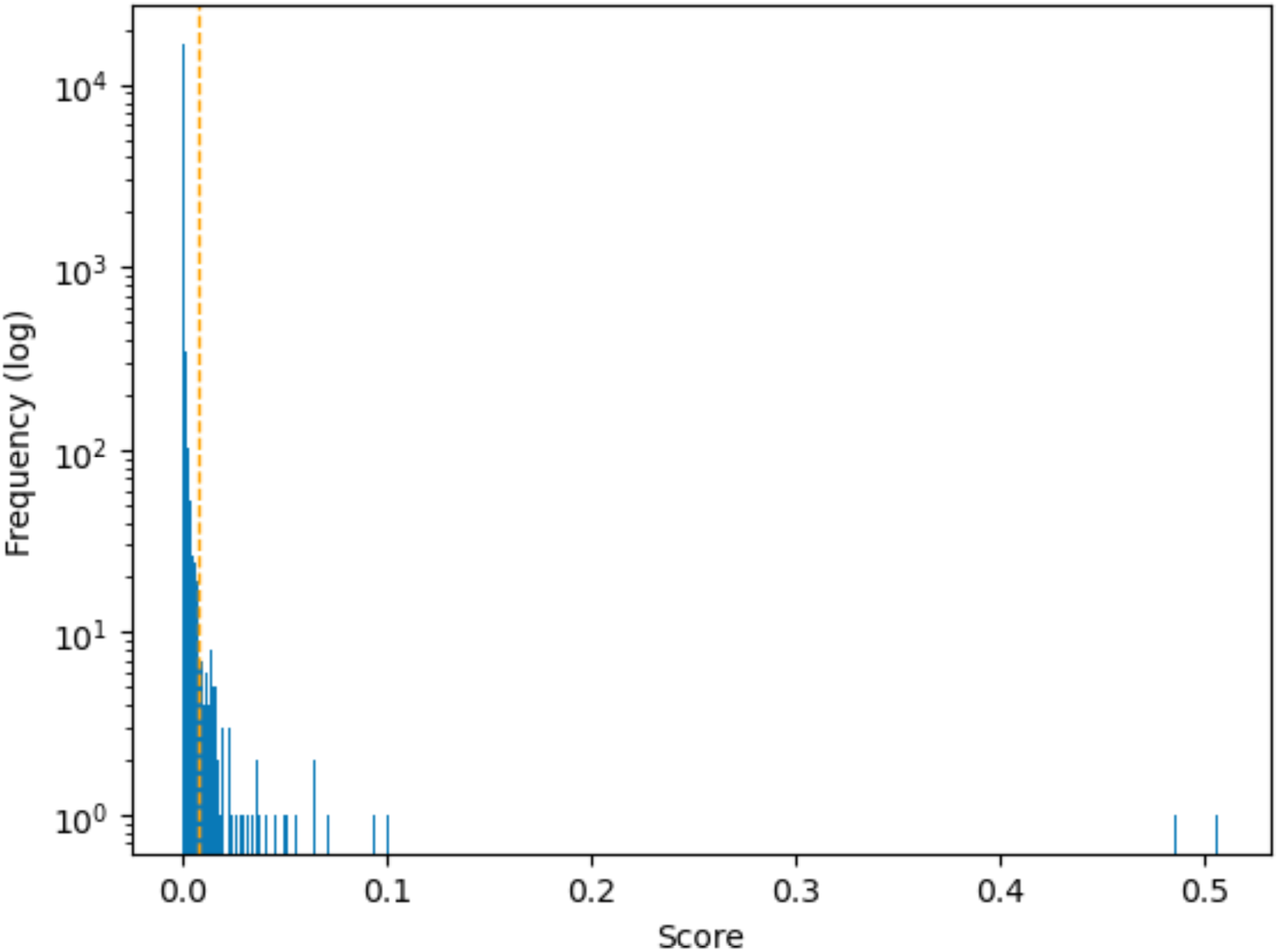
The distribution of scores for the knockouts. To suggest candidate SL pairs, we utilize a score cutoff of 0.0071 (shown as a dashed vertical line). This results in 92 knockouts.

**SI Figure 5:**
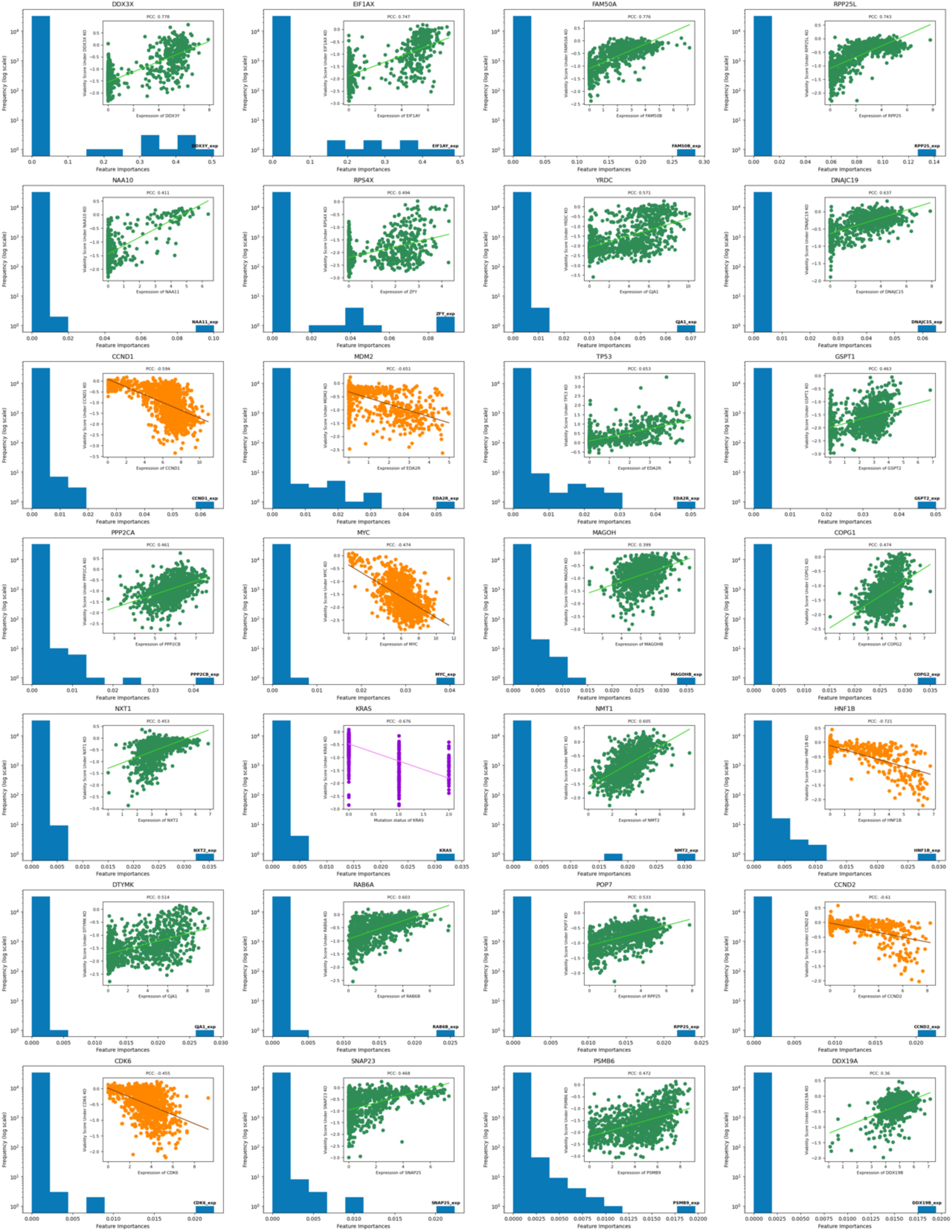
SL-RFM feature importance plots 1-28 of our proposed SL pairs.

**SI Figure 6:**
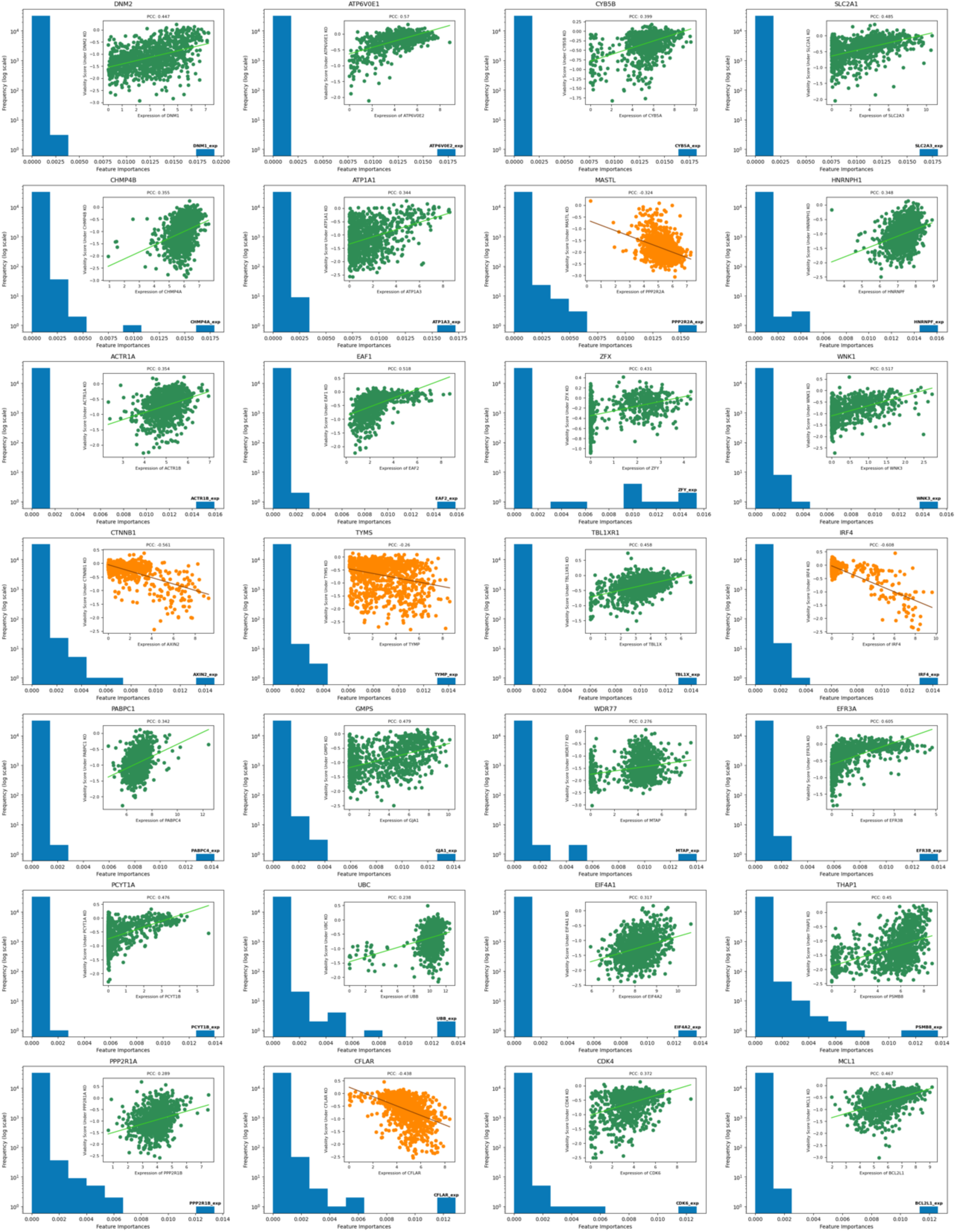
SL-RFM feature importance plots 29-56 of our proposed SL pairs.

**SI Figure 7:**
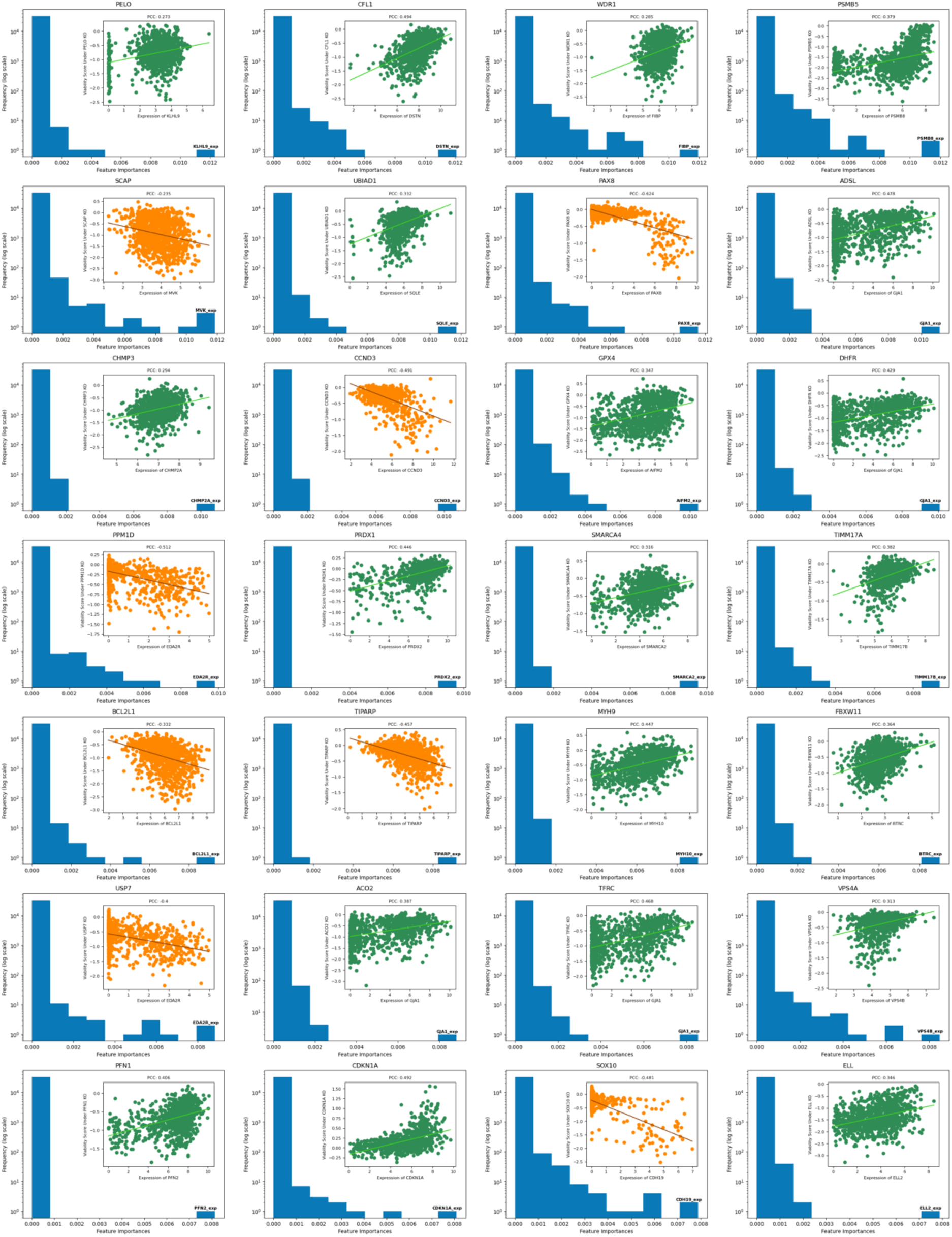
SL-RFM feature importance plots 57-84 of our proposed SL pairs.

**SI Figure 8:**
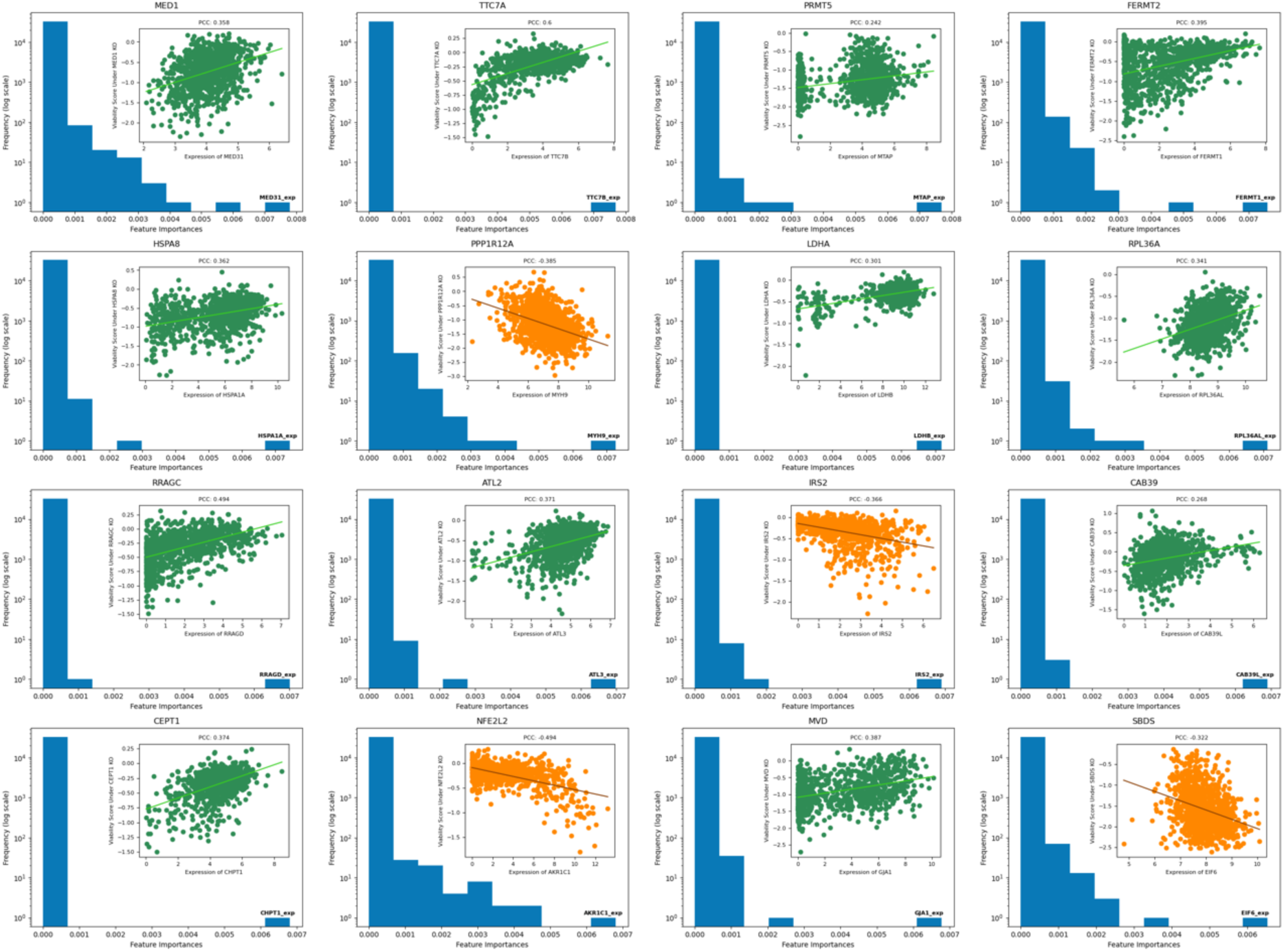
SL-RFM feature importance plots 85-100 of our proposed SL pairs.

**SI Figure 9:**
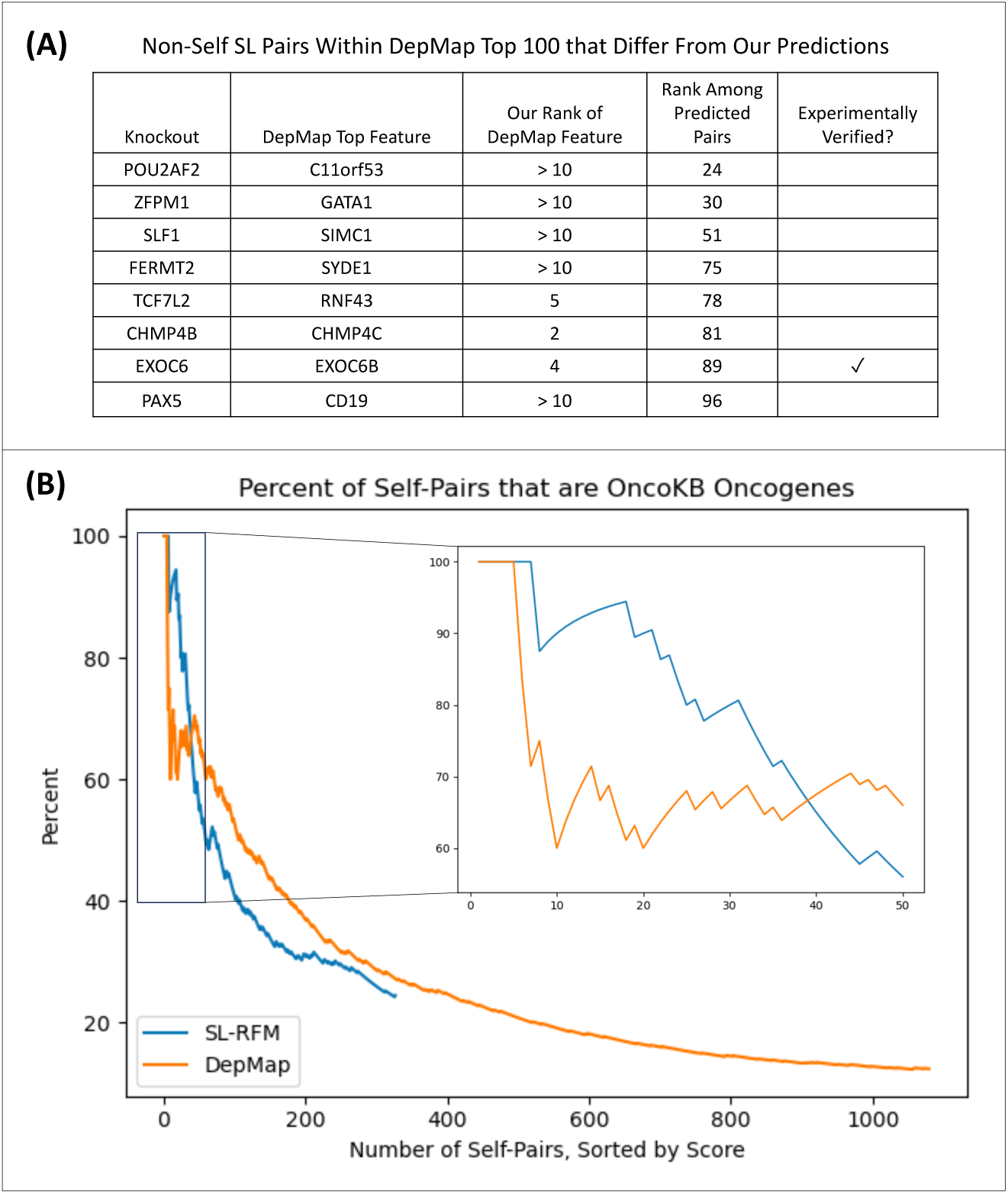
Comparison of the SL pairs suggested by our pipeline to those suggested by DepMap. **(A)** A list of the non-self SL pairs out of the top 100 pairs proposed by the best model from DepMap that differ from pairs proposed by our pipeline. We sort DepMap pairs by score, as defined by difference between max feature importance and mean feature importance for a given knockout. Since DepMap only published the feature importances of the top 10 features for each knockout, the score was calculated using the 10 provided features. Note that only two out of ten of the top pairs of DepMap not found using our method are experimentally verified. **(B)** Comparison of the percent of self-pairs proposed by our method and DepMap that are verified oncogenes in OncoKB. We observe that the model from DepMap proposes many more self-pairs than our method, but that most of these pairs are not verified oncogenes in OncoKB.

**SI Figure 10:**
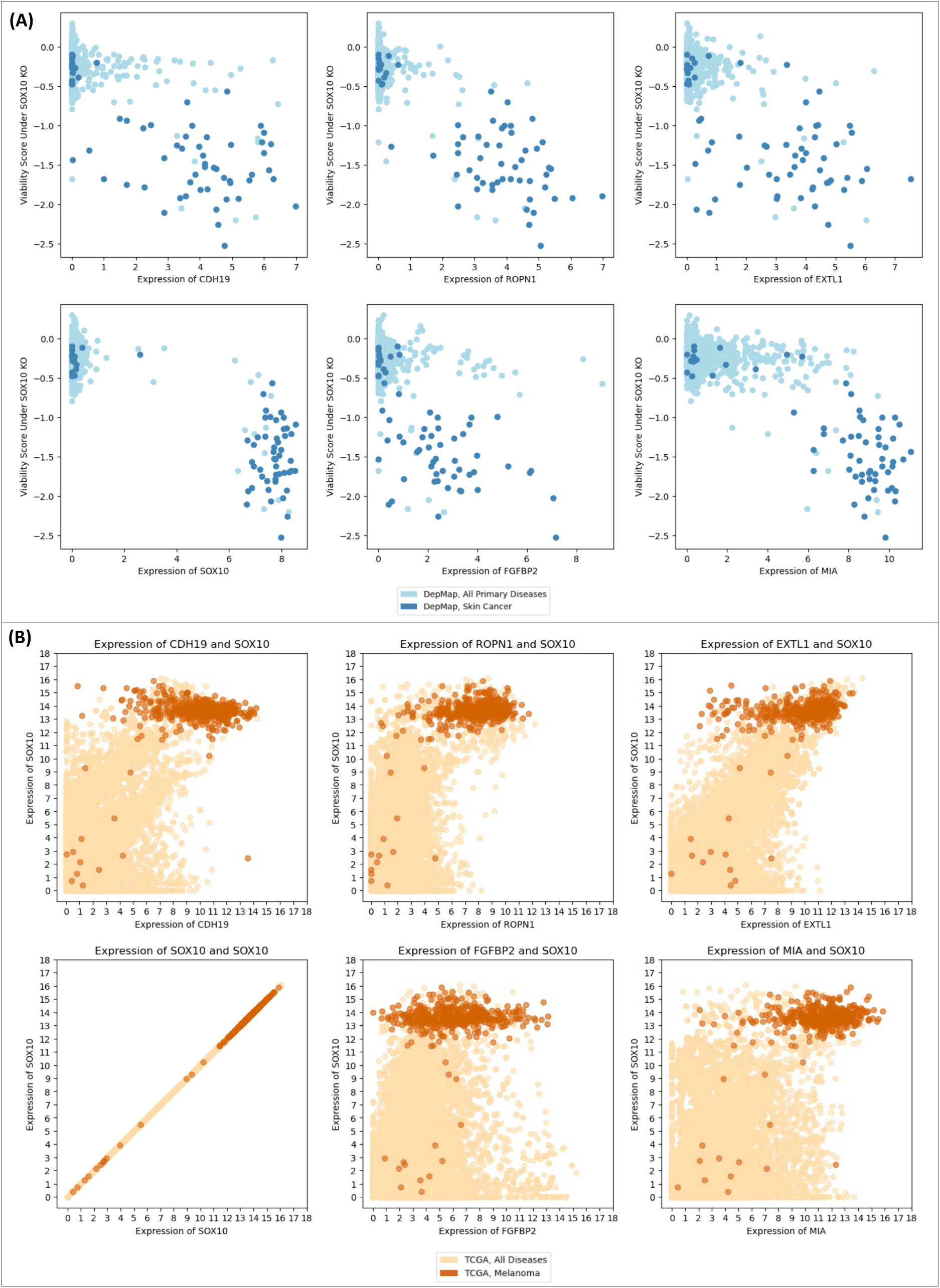
**(A)** A visualization of the relationship between expression of genes associated with the top six most important features for predicting viability scores under knockout of SOX10 using SL-RFM. The majority of cell lines with over-expression of the targets are derived from skin cancer and have low viability under the SOX10 knockout. **(B)** Joint expression of these genes and SOX10 in TCGA. The majority of melanoma samples have SOX10 overexpressed. These analyses suggest that knocking out SOX10 causes low viability for cells with over-expression of the genes associated with these top features, which occurs frequently in melanoma.

**SI Figure 11:**
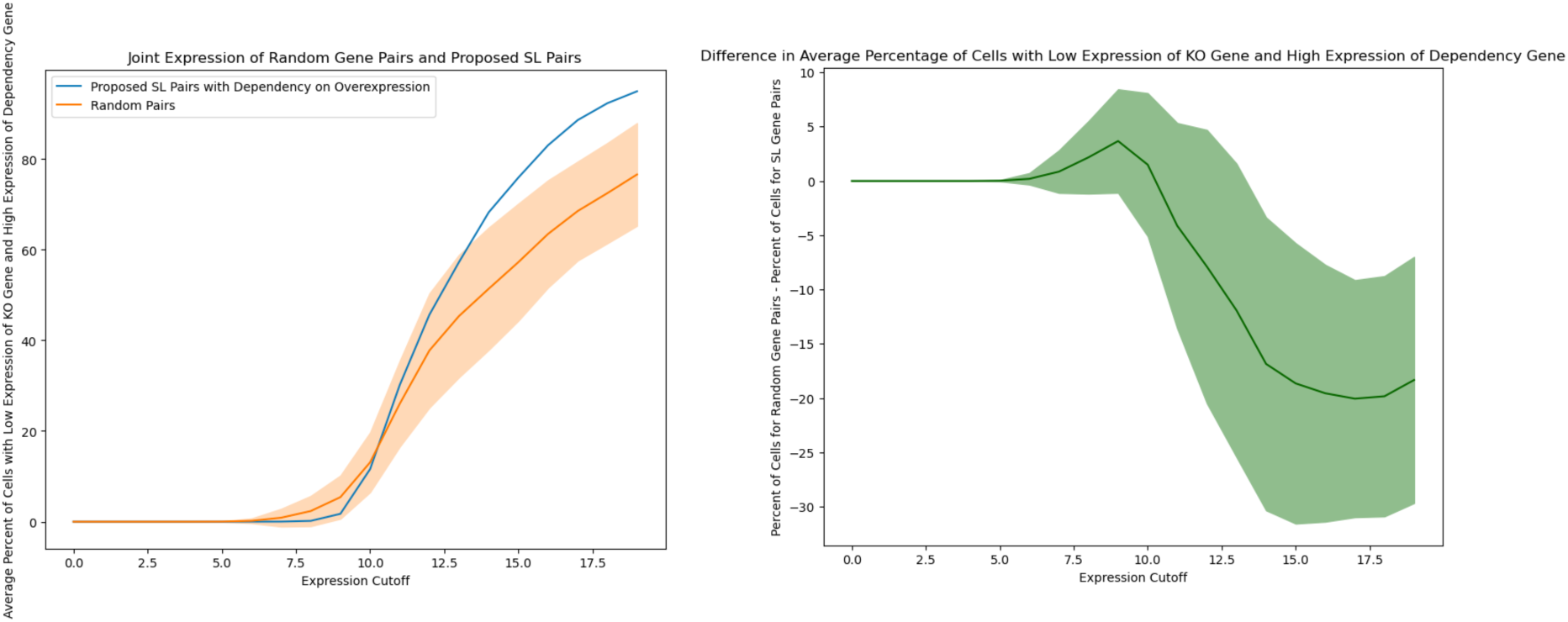
Visualization showing that for SL pairs of the form (A, B) with dependency on knockout of gene A and over-expression of gene B, there are few TCGA samples exhibiting low expression of gene A and high expression of gene B. For 9 randomly sampled gene pairs (sampled 10 times) and the 9 SL pairs with dependency on over-expression of the form (A, B), we plot the average percentage of TCGA samples that have expression of the gene A below the expression cutoff, *c*, and the expression of the gene B above 20 − *c*. In the figure on the right, we plot the difference between the two curves. For *c <* 10, there are up to 10% fewer TCGA samples with low expression of gene A and high expression of gene B than random samples. As *c* increases, this relationship flips since there can be many samples for which gene B is under-expressed.

**SI Figure 12:**
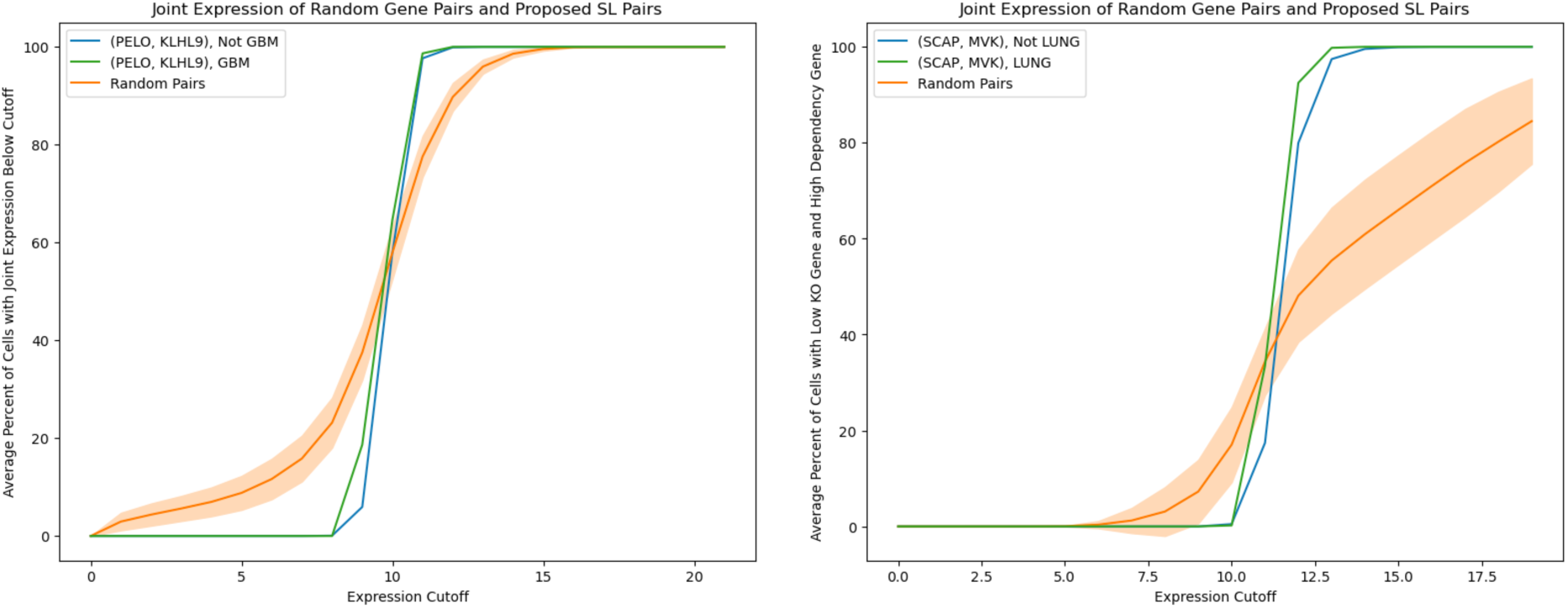
Visualization showing that for the SL pairs we propose, there are fewer TCGA samples with expression patterns corresponding to proposed synthetically lethal gene interactions than random pairs. The dependency of PELO/KLHL9 is on under-expression, so we validate that there are fewer TCGA samples with the under-expression of both genes than random pairs. The dependencies of SCAP/MVK and SOX10/CDH19 are on over-expression, so we validate that for expression cutoff below 10, there are fewer TCGA samples with the under-expression of SCAP/SOX10 and over-expression of MVK/CDH19. For SCAP/MVK and SOX10/CDH19, we plot the average percentage of TCGA samples that have expression of SCAP/SOX10 below the expression cutoff, *c*, and the expression of MVK/CDH19 above 20−*c*. We also plot each of these curves upon stratifying the TCGA samples by cancer type based on our analysis in Fig. 4. Namely, the susceptible cancer type for PELO/KLHL9 is glioblastoma (GBM), the type for SCAP/MVK is lung cancer (LUNG), and the type for SOX10/CDH19 is skin cancer (SCKM).

**SI Figure 13:**
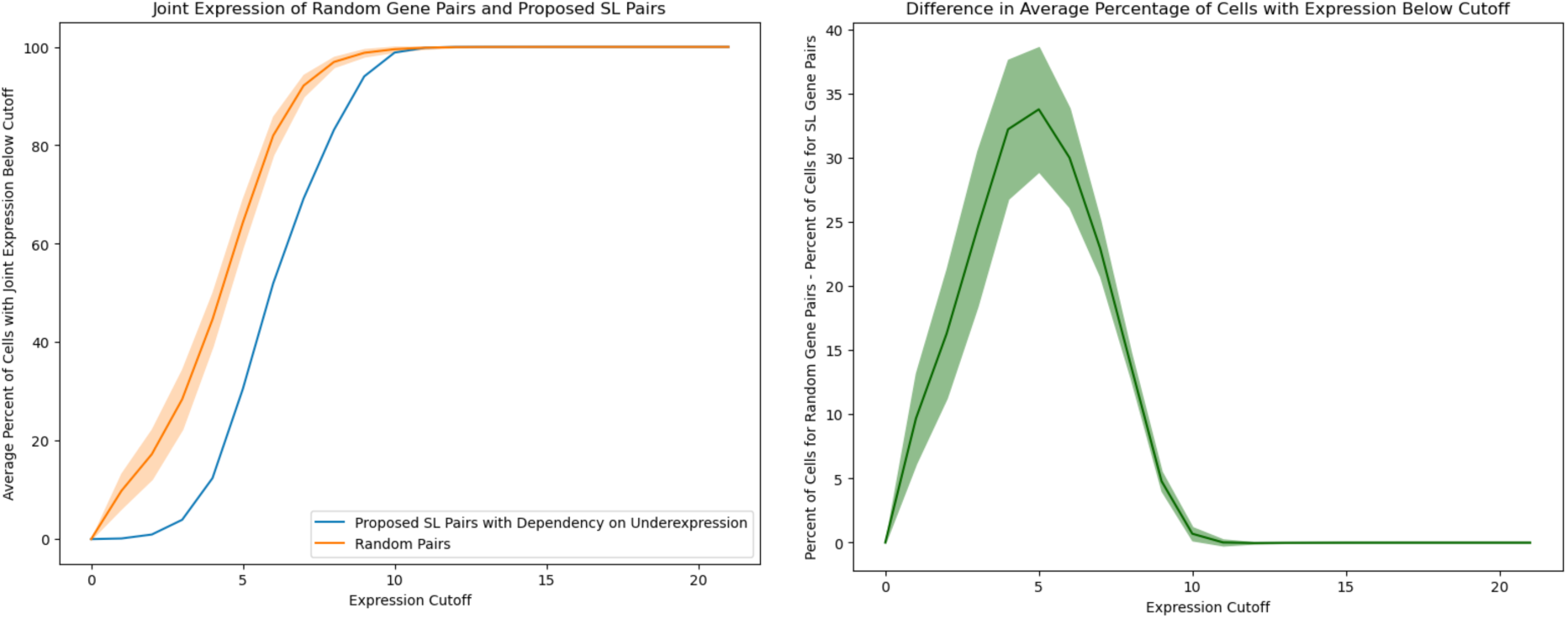
Visualization showing that SL genes with a dependency on under-expression are not simultaneously under-expressed in GTEx data. The GTEx TPM expression data was normalized as log_2_(TPM + 1). For 67 randomly sampled gene pairs (sampled 10 times) and the 67 out of 92 SL pairs with a dependency on under-expression and with data in TCGA (we omitted RPP25L/RPP25, COPG1/COPG2, and CHMP3/CHMP2A for this reason), we plot the percentage of GTEx samples for which both genes have an expression below the given x-axis coordinate. On the right, we plot the difference of the percentage of GTEx samples with expression of randomly sampled gene pairs below the cutoff and the percentage of GTEx samples with expression of SL pairs with expression below the cutoff. Overall, we observe up to a 40% difference, indicating that our candidate pairs are almost never simultaneously under-expressed in GTEx samples.

