## Supporting Information for "Synthetic Lethality Screening with Recursive Feature Machines"

**(A)**

Comparing Performance in Predicting Cell Viability, All Genes

| | Laplacian Kernel | Gaussian Kernel | Mean Over Cell Type |
|---|---|---|---|
| Pearson R | **0.935** | 0.933 | 0.929 |
| $R^2$ | **0.875 ± 0.040** | 0.872 ± 0.041 | 0.865 ± 0.041 |
| Cosine Similarity | **0.943 ± 0.019** | 0.942 ± 0.020 | 0.938 ± 0.020 |

Comparing Performance in Predicting Cell Viability, SL Genes

| | Laplacian Kernel | Gaussian Kernel | Mean Over Cell Type |
|---|---|---|---|
| Pearson R | 0.742 | **0.744** | 0.664 |
| $R^2$ | **0.522 ± 0.209** | 0.522 ± 0.216 | 0.413 ± 0.227 |
| Cosine Similarity | 0.840 ± 0.072 | **0.841 ± 0.072** | 0.797 ± 0.083 |

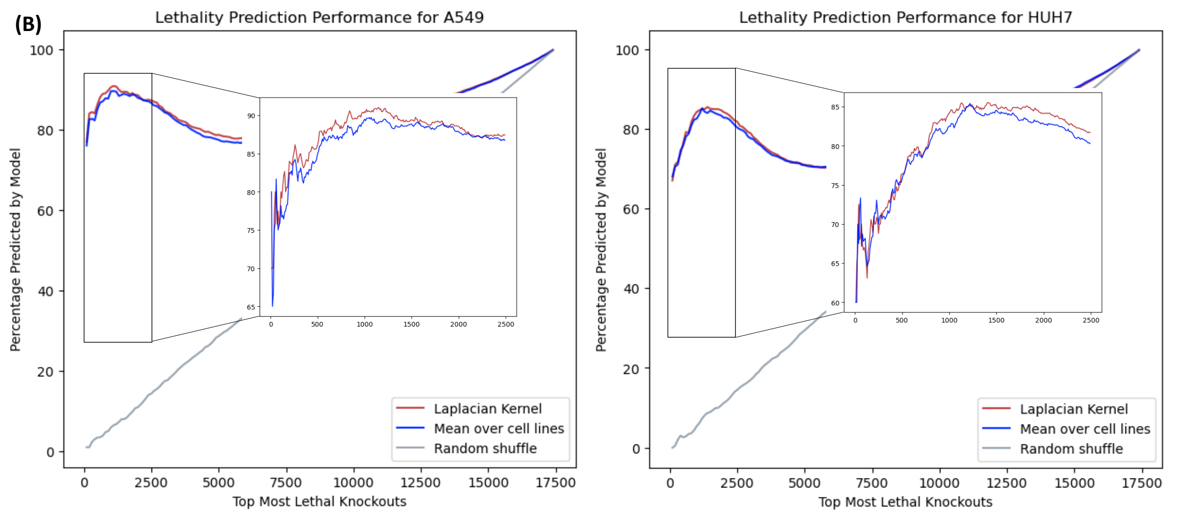


SI Figure 1: Comparison of model performance in predicting viability data. All metrics are described in Methods. **(A)** Comparison of RFM with varying base kernels (Laplace and Gaussian kernel) on predicting cell viability, where test cell lines are held out using 5-fold cross validation. **(B)** Analyzing predictions for individual cell lines from five-fold cross-validation. We compare the list of predicted knockouts and the list of ground truth knockouts sorted by viability score for sample cell lines (A549 and HUH7). We observe that RFM with the Laplace kernel base predictor outperforms both a mean over cell line benchmark and a random baseline, illustrating the effectiveness of our selected model on held-out test data.

Effectiveness of Our Pipeline

| Experimentally Verified SL Pairs from [1] | PCC | PCC | PARIS | PARIS | DepMap | DepMap | **SL-RFM (Ours)** | **SL-RFM (Ours)** |
|---|---|---|---|---|---|---|---|---|
| SMARCA2/SMARCA4 | **1** | **1** | **1** | 647 | **1** | **1** | **1** | **1** |
| ARID1B/ARID1A | **1** | 7837 | **1** | 419 | **1** | > 10 | **1** | 228 |
| STAG1/STAG2 | **1** | **1** | **1** | 3133 | **1** | **1** | **1** | **1** |
| CREBBP/EP300 | **1** | 5077 | **1** | 435 | **1** | > 10 | **1** | 14578 |
| VPS4A/VPS4B | 3 | 10 | 3471 | 4150 | 5 | 5 | **1** | **1** |
| DDX17/DDX5 | **1** | **1** | **1** | 1071 | **1** | **1** | **1** | **1** |
| ENO1/ENO2 | 4 | 1690 | **1** | 5070 | **1** | > 10 | **1** | 8378 |
| SMARCC2/SMARCC1 | 6 | 822 | **1** | 1577 | **1** | > 10 | **1** | 1791 |
| UBC/UBB | 2 | N/A | **1** | N/A | > 10 | > 10 | **1** | N/A |
| MAGOH/MAGOHB | **1** | 15 | **1** | 5200 | **1** | > 10 | **1** | 2 |
| ME2/ME3 | 14850 | 15640 | 3307 | 4944 | > 10 | > 10 | 15938 | 16959 |
| FAM50A/FAM50B | **1** | 3 | **1** | 530 | **1** | **1** | **1** | **1** |

SI Figure 2: Ranks of all models from Fig. 2 when considering the first gene in the pair as the knockout (white) or the second gene in the pair as the knockout (gray). The minimum rank between white and gray columns is reported in Fig. 2. We note that DepMap only provides the top 10 most important features for prediction. If the SL pair gene does not appear among the top 10, we denote the rank as $> 10$. All other models return the full set of feature importances.

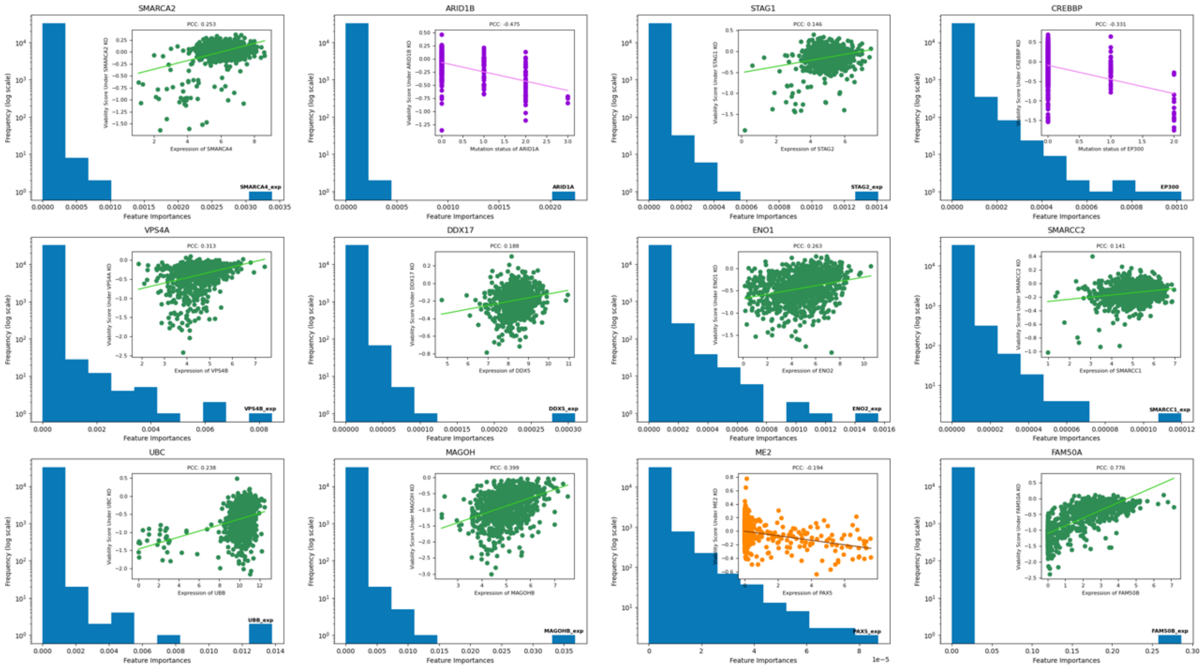


SI Figure 3: SL-RFM feature importance plots for knockouts part of SL pairs from [18].

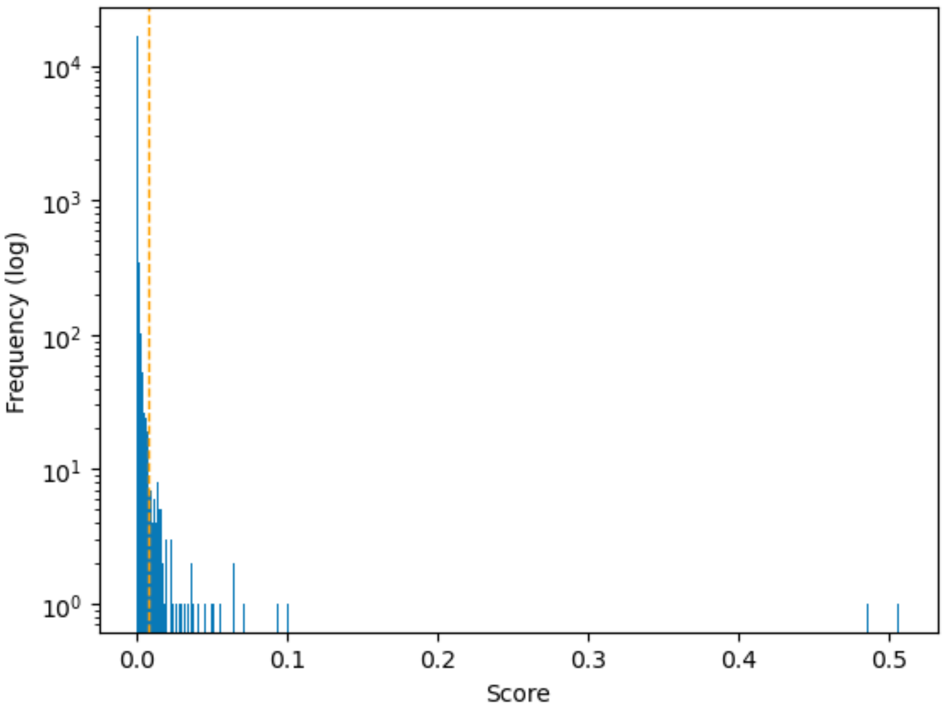


SI Figure 4: The distribution of scores for the knockouts. To suggest candidate SL pairs, we utilize a score cutoff of 0.0071 (shown as a dashed vertical line). This results in 92 knockouts.

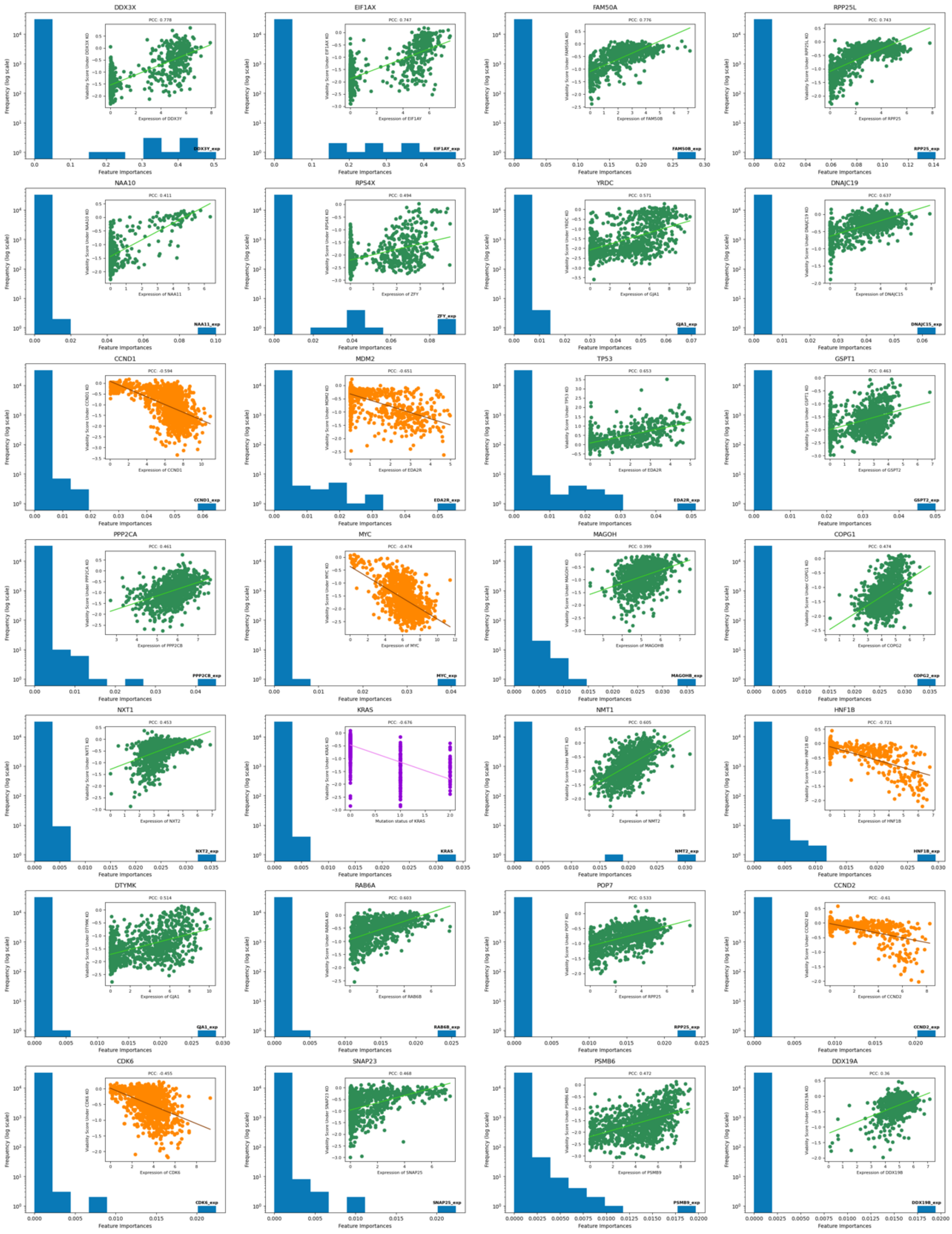


SI Figure 5: SL-RFM feature importance plots 1-28 of our proposed SL pairs.

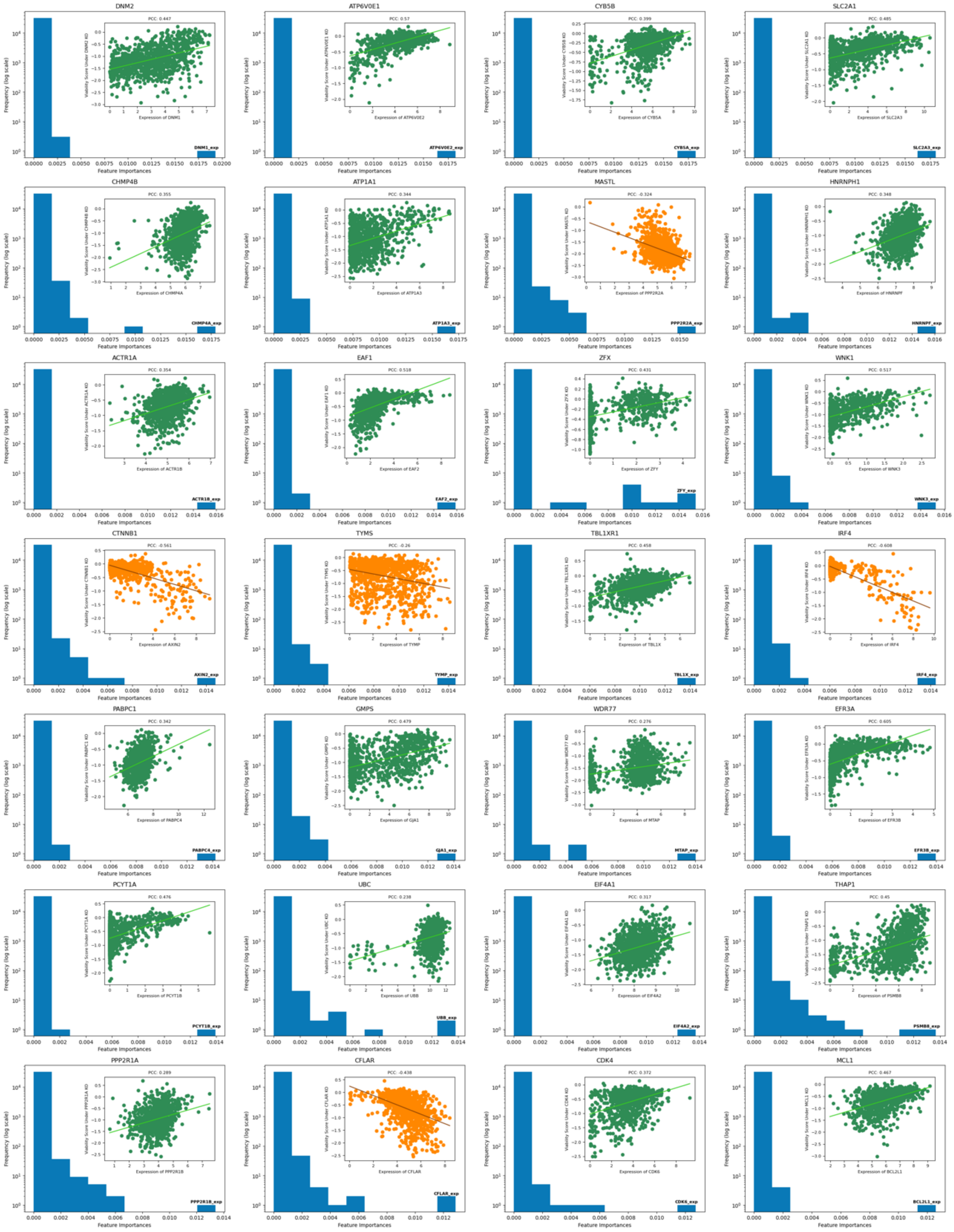


SI Figure 6: SL-RFM feature importance plots 29-56 of our proposed SL pairs.

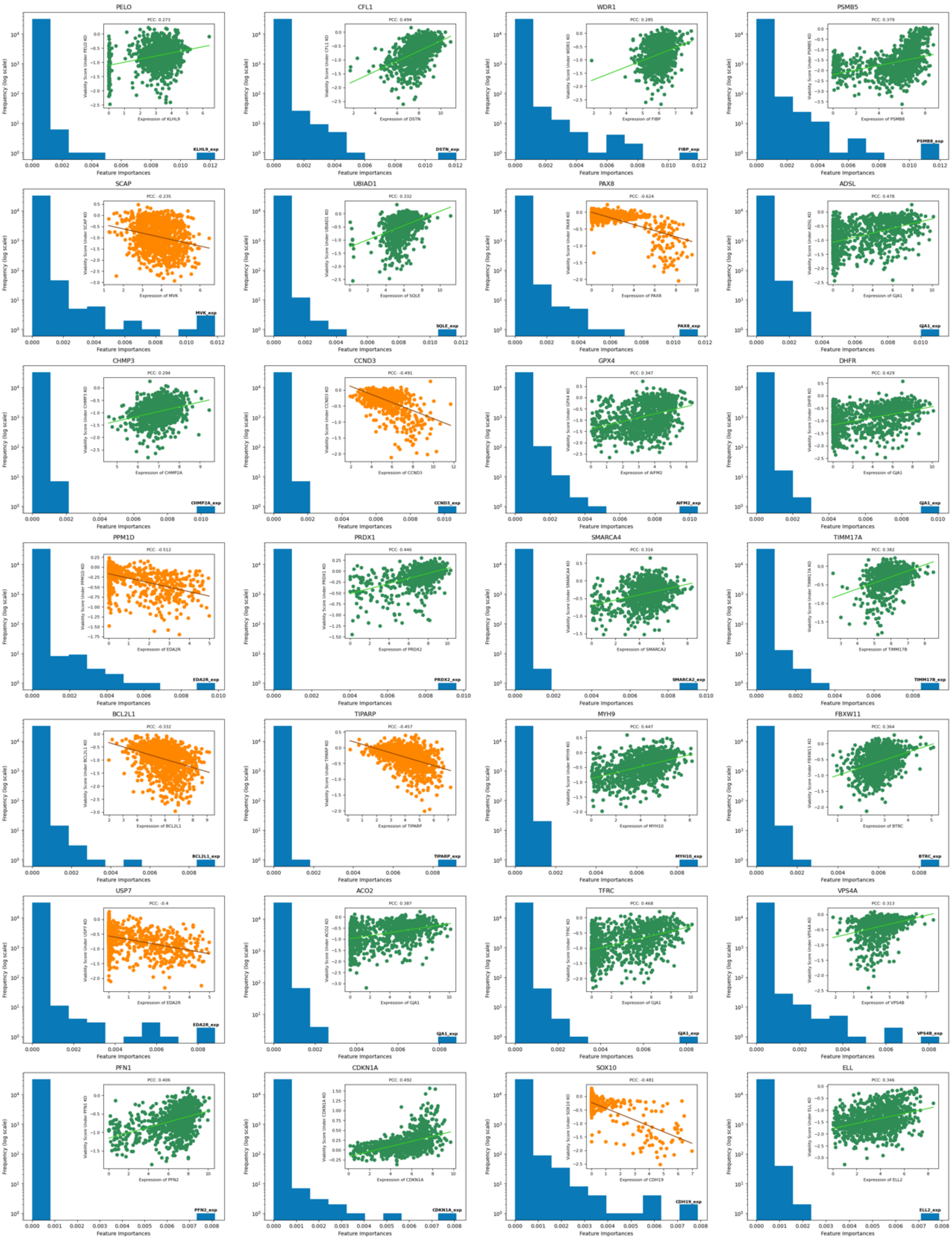


SI Figure 7: SL-RFM feature importance plots 57-84 of our proposed SL pairs.

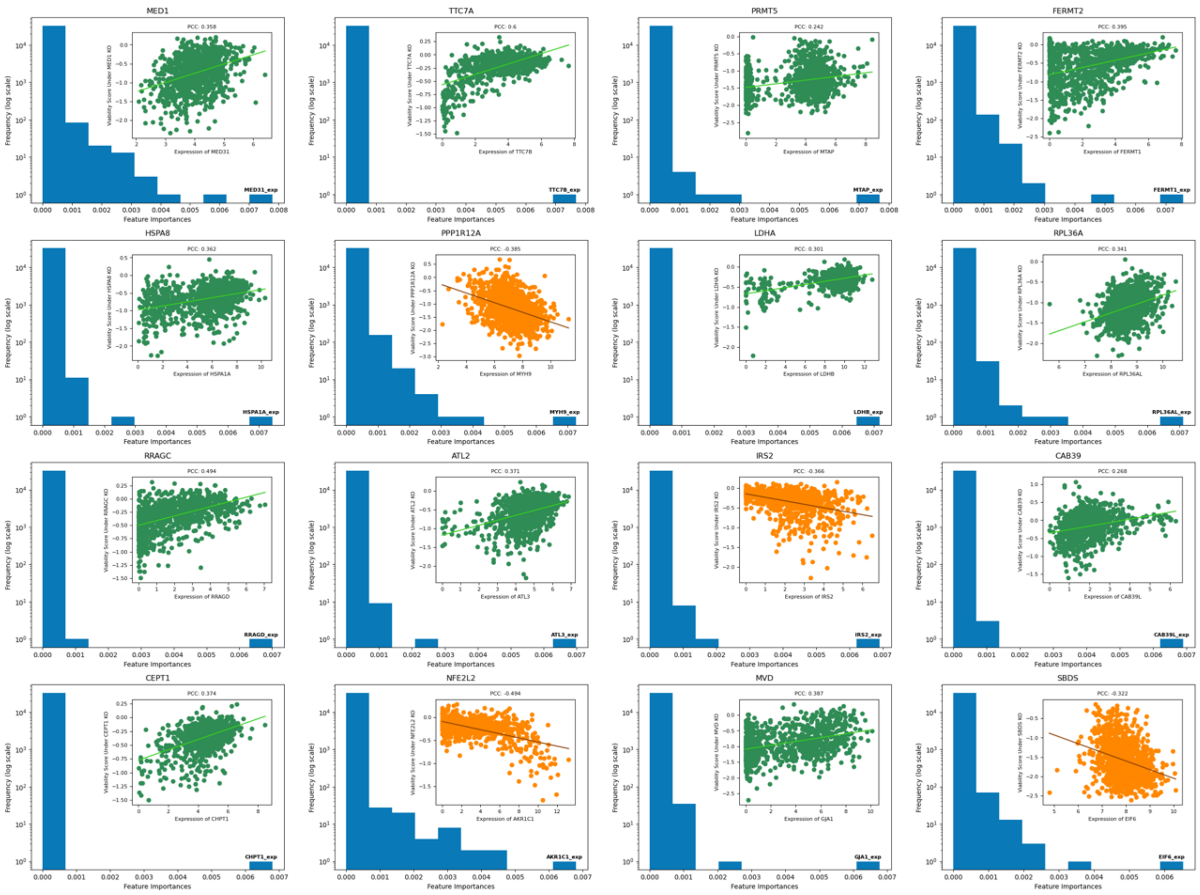


SI Figure 8: SL-RFM feature importance plots 85-100 of our proposed SL pairs.

**(A)** Non-Self SL Pairs Within DepMap Top 100 that Differ From Our Predictions

| Knockout | DepMap Top Feature | Our Rank of DepMap Feature | Rank Among Predicted Pairs | Experimentally Verified? |
|---|---|---|---|---|
| POU2AF2 | C11orf53 | > 10 | 24 | |
| ZFPM1 | GATA1 | > 10 | 30 | |
| SLF1 | SIMC1 | > 10 | 51 | |
| FERMT2 | SYDE1 | > 10 | 75 | |
| TCF7L2 | RNF43 | 5 | 78 | |
| CHMP4B | CHMP4C | 2 | 81 | |
| EXOC6 | EXOC6B | 4 | 89 | ✓ |
| PAX5 | CD19 | > 10 | 96 | |

**(B)**

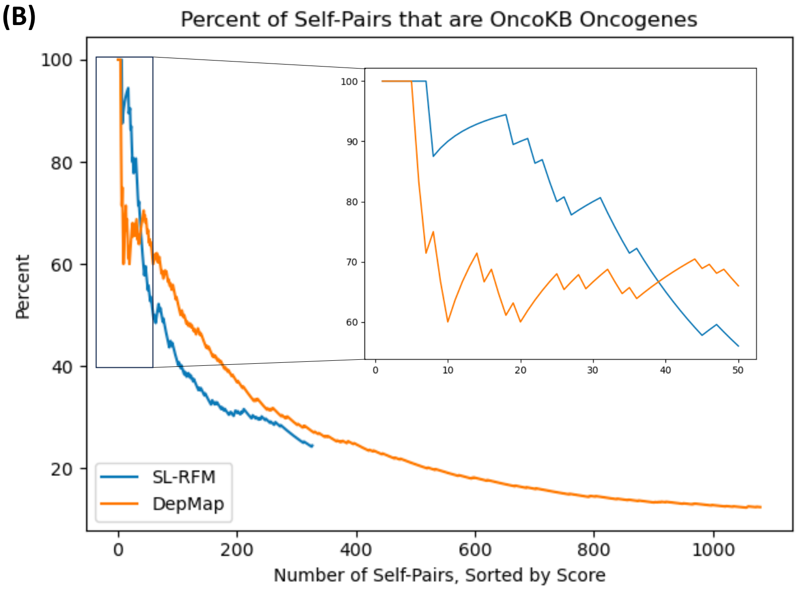


SI Figure 9: Comparison of the SL pairs suggested by our pipeline to those suggested by DepMap. **(A)** A list of the non-self SL pairs out of the top 100 pairs proposed by the best model from DepMap that differ from pairs proposed by our pipeline. We sort DepMap pairs by score, as defined by difference between max feature importance and mean feature importance for a given knockout. Since DepMap only published the feature importances of the top 10 features for each knockout, the score was calculated using the 10 provided features. Note that only two out of ten of the top pairs of DepMap not found using our method are experimentally verified. **(B)** Comparison of the percent of self-pairs proposed by our method and DepMap that are verified oncogenes in OncoKB. We observe that the model from DepMap proposes many more self-pairs than our method, but that most of these pairs are not verified oncogenes in OncoKB.

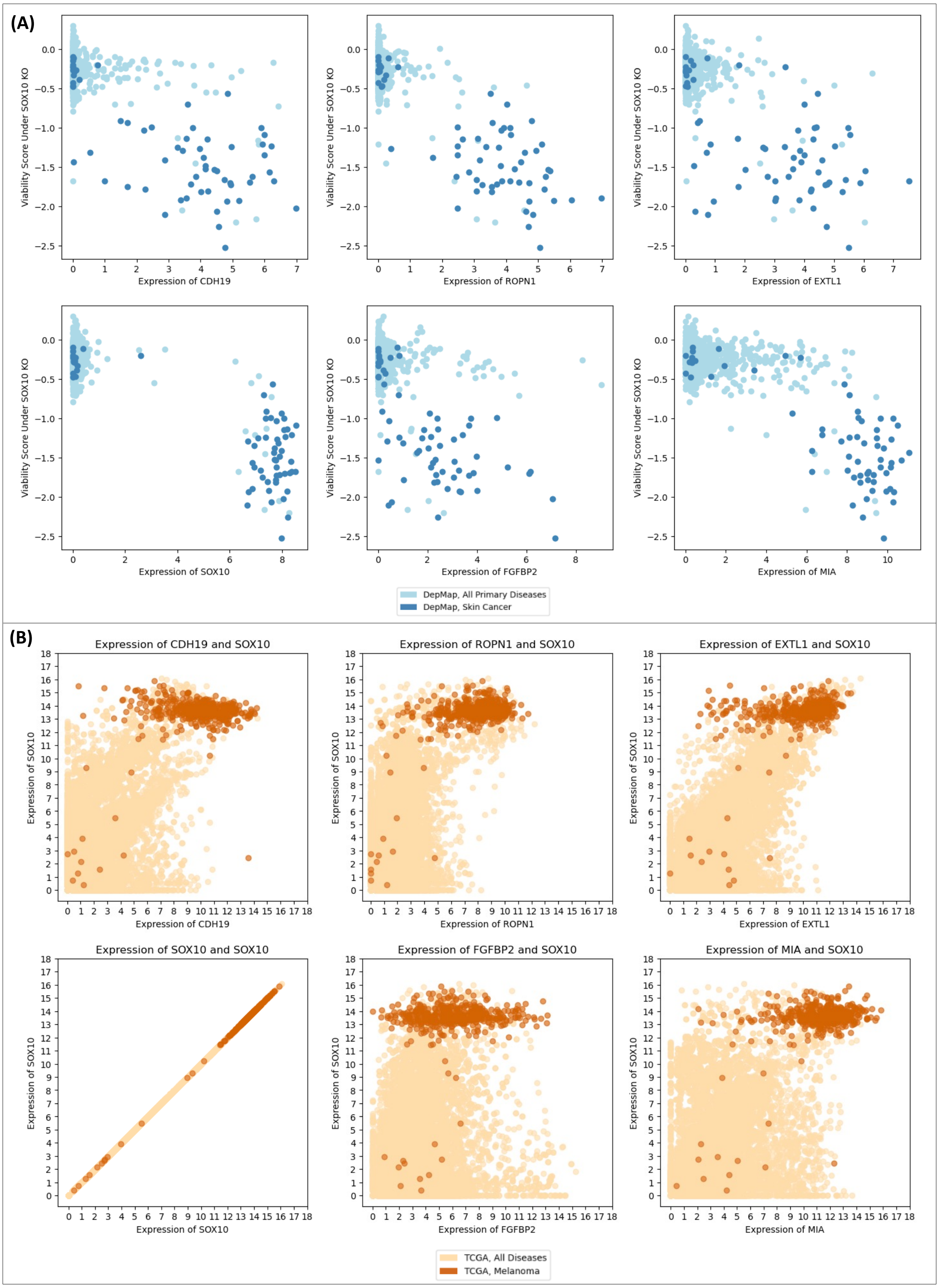


SI Figure 10: **(A)** A visualization of the relationship between expression of genes associated with the top six most important features for predicting viability scores under knockout of SOX10 using SL-RFM. The majority of cell lines with over-expression of the targets are derived from skin cancer and have low viability under the SOX10 knockout. **(B)** Joint expression of these genes and SOX10 in TCGA. The majority of melanoma samples have SOX10 overexpressed. These analyses suggest that knocking out SOX10 causes low viability for cells with over-expression of the genes associated with these top features, which occurs frequently in melanoma.

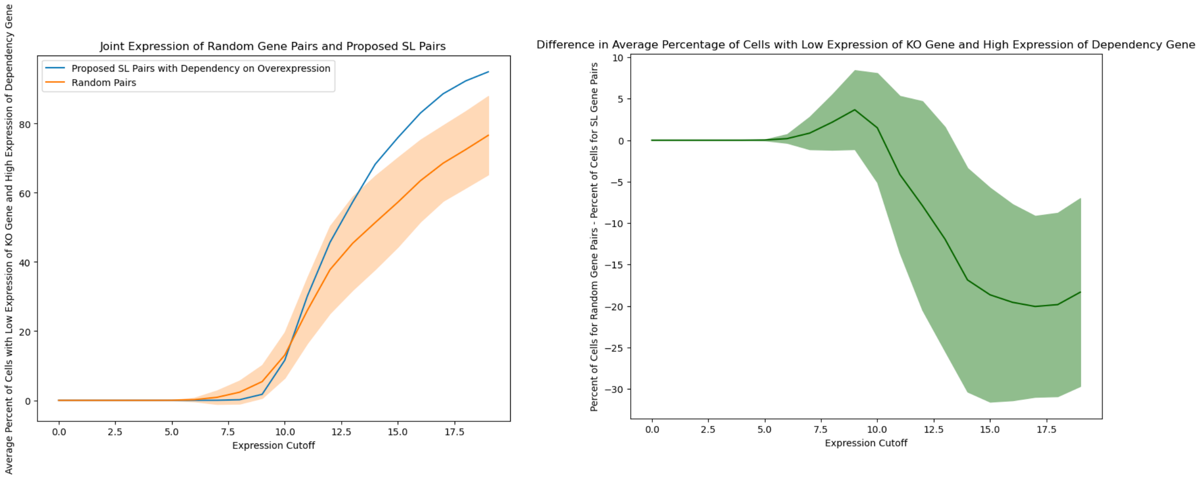


SI Figure 11: Visualization showing that for SL pairs of the form (A, B) with dependency on knockout of gene A and over-expression of gene B, there are few TCGA samples exhibiting low expression of gene A and high expression of gene B. For 9 randomly sampled gene pairs (sampled 10 times) and the 9 SL pairs with dependency on over-expression of the form (A, B), we plot the average percentage of TCGA samples that have expression of the gene A below the expression cutoff, $c$, and the expression of the gene B above $20 - c$. In the figure on the right, we plot the difference between the two curves. For $c < 10$, there are up to 10% fewer TCGA samples with low expression of gene A and high expression of gene B than random samples. As $c$ increases, this relationship flips since there can be many samples for which gene B is under-expressed.

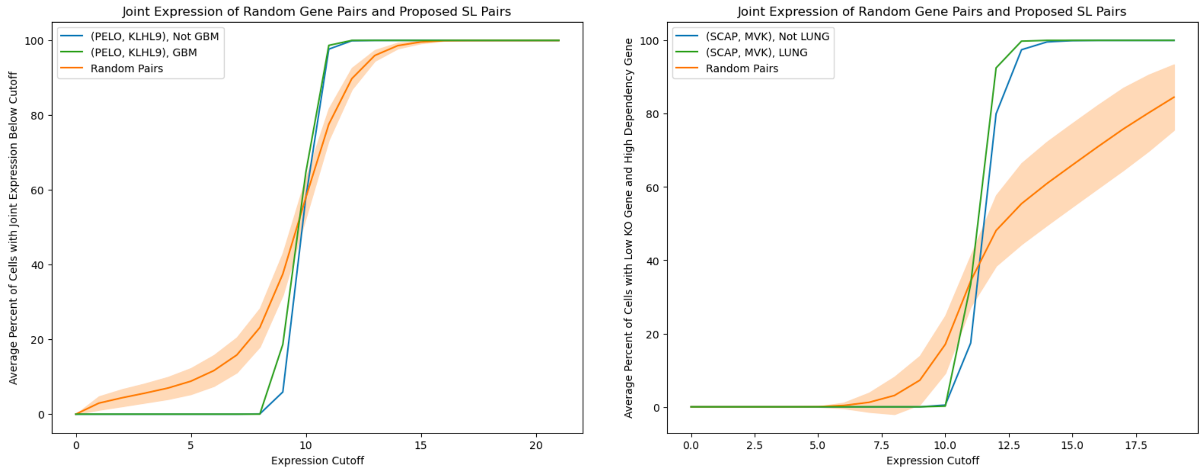


SI Figure 12: Visualization showing that for the SL pairs we propose, there are fewer TCGA samples with expression patterns corresponding to proposed synthetically lethal gene interactions than random pairs. The dependency of PELO/KLHL9 is on under-expression, so we validate that there are fewer TCGA samples with the under-expression of both genes than random pairs. The dependencies of SCAP/MVK and SOX10/CDH19 are on over-expression, so we validate that for expression cutoff below 10, there are fewer TCGA samples with the under-expression of SCAP/SOX10 and over-expression of MVK/CDH19. For SCAP/MVK and SOX10/CDH19, we plot the average percentage of TCGA samples that have expression of SCAP/SOX10 below the expression cutoff, $c$, and the expression of MVK/CDH19 above $20-c$. We also plot each of these curves upon stratifying the TCGA samples by cancer type based on our analysis in Fig. 4. Namely, the susceptible cancer type for PELO/KLHL9 is glioblastoma (GBM), the type for SCAP/MVK is lung cancer (LUNG), and the type for SOX10/CDH19 is skin cancer (SCKM).

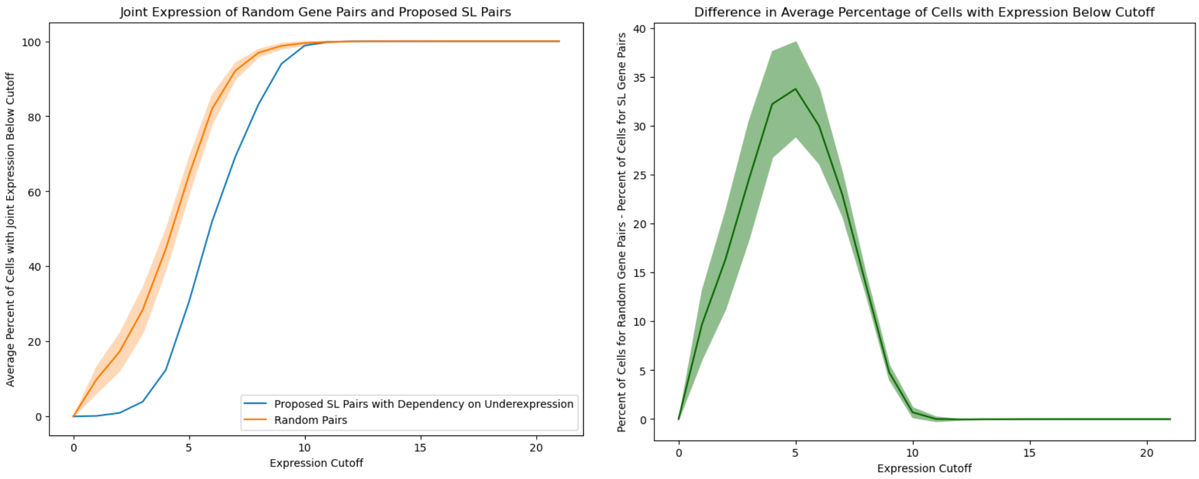


SI Figure 13: Visualization showing that SL genes with a dependency on under-expression are not simultaneously under-expressed in GTEx data. The GTEx TPM expression data was normalized as $\log_2(\text{TPM} + 1)$. For 67 randomly sampled gene pairs (sampled 10 times) and the 67 out of 92 SL pairs with a dependency on under-expression and with data in TCGA (we omitted RPP25L/RPP25, COPG1/COPG2, and CHMP3/CHMP2A for this reason), we plot the percentage of GTEx samples for which both genes have an expression below the given x-axis coordinate. On the right, we plot the difference of the percentage of GTEx samples with expression of randomly sampled gene pairs below the cutoff and the percentage of GTEx samples with expression of SL pairs with expression below the cutoff. Overall, we observe up to a 40% difference, indicating that our candidate pairs are almost never simultaneously under-expressed in GTEx samples.
